# *α*-actinin-4 drives invasiveness by regulating myosin IIB expression and myosin IIA localization

**DOI:** 10.1101/2021.01.12.426368

**Authors:** Amlan Barai, Abhishek Mukherjee, Alakesh Das, Neha Saxena, Shamik Sen

**Affiliations:** Dept. of Biosciences & Bioengineering, IIT Bombay, Mumbai, India; IITB-Monash Research Academy, Mumbai, India; Dept. of Mechanical Engineering, IIT Bombay, Mumbai, India; Dept. of Biological Regulation, Weizmann Institute of Science, Israel

## Abstract

The mechanisms by which the mechanoresponsive actin crosslinking protein *α*-actinin-4 (ACTN4) regulates cell motility and invasiveness remains incompletely understood. Here we show that in addition to regulating protrusion dynamics and focal adhesion formation, ACTN4 transcriptionally regulates expression of non-muscle myosin IIB (NMM IIB), which is essential for mediating nuclear translocation during 3D invasion. We further demonstrate association between NMM IIA and ACTN4 at the cell front ensures retention of NMM IIA at the cell periphery. A protrusion-dependent model of confined migration recapitulating experimental observations predicts a dependence of protrusion forces on the degree of confinement and on the ratio of nucleus to matrix stiffness. Together, our results suggest that ACTN4 is a master regulator of cancer invasion that regulates invasiveness by controlling NMM IIB expression and NMM IIA localization.

## Introduction

Cancer metastasis, i.e., spread of cancer cells from one tissue to another, is the primary reason behind the high mortality in cancer. Consequently, identifying molecules mediating cancer metastasis and their mechanisms of action can provide us with useful strategies for therapeutic intervention. Most epithelial cancers are marked by dramatic alterations in composition and organization of the extracellular matrix (ECM)^1,2^. ECM stiffening, induced by excessive deposition of collagen I and its crosslinking by lysyl oxidase reported in breast cancer, has been shown to be capable of driving cancer progression^3,4^. For successful invasion through these dense matrices, cancer cells are known to upregulate expression of matrix degrading enzymes such as matrix metalloproteinases (MMPs) for locally degrading the matrix and creating paths for escape^5–7^. However, even after such localized degradation, translocation of the nucleus, which is large and stiff, represents a rate limiting factor for confined migration^8^. Nuclear translocation is mediated through a combination of non-muscle myosin IIA (NMM IIA)-based pulling from the cell front and NMM IIB-based squeezing of the nucleus from the rear ^9^.

*α*-actinin-4 (ACTN4) is a non-muscle actin crosslinking protein whose expression has been strongly correlated with cancer invasiveness, metastasis and therapeutic resistance across different types of cancers^10–14^. These behaviors are attributed to involvement of ACTN4 with several well-known signaling molecules including *β*-catenin^15^, NFkB^16^ and AKT^14^. Recently, ACTN4 along with filamin A, non-muscle myosin IIA and IIB were identified as four mechanoresponsive proteins associated with the actin cytoskeleton which were enriched at sites of localized pressure^17^. While these studies illustrate the importance of ACTN4 as a central player regulating several aspects of cancer invasion, the mechanisms by which it contributes to increased invasiveness remains incompletely understood.

In this study, we have specifically probed the role of ACTN4 in regulating cancer invasiveness. Using TCGA^18–20^ database and breast cancer cell lines of varying invasiveness, we first establish a positive correlation between ACTN4 levels and cell motility. We then use high ACTN4 expressing MDA-MB-231 breast cancer cells and HT-1080 fibrosarcoma cells to probe the functions of ACTN4 in regulating cell migration and invasion. We show that ACTN4 knockdown leads to cell elongation, formation of smaller focal adhesions, but reduction in protrusion-retraction dynamics. Using collagen gel invasion and transwell pore migration experiments, we then show that ACTN4 regulates nuclear translocation by modulating NMM IIB expression. We further show that ACTN4 directly associates with NMM IIA at the cell front, with this association essential for peripheral localization of NMM IIA. Finally, using a computational model, we estimate ACTN4-mediated protrusive forces at the cell front required for confined migration. Collectively, our results suggest that ACTN4 regulates cancer invasiveness by regulating nuclear deformation via modulation of NMM IIB expression and by stabilizing protrusions at the cell front via its association with NMM IIA.

## Materials and methods

### Experiments

#### Cell culture

MCF-7, T-47D, ZR-75-1, MDA-MB-231, and HT-1080 cells were obtained from National Center for Cell Science (NCCS, Pune, India) with MCF-7 cultured in MEM (Gibco, Cat #61100-061), T-47D and ZR-75-1 were cultured in RPMI-1640 (Gibco, Ref# 31800-022), and MDA-MB-231 and HT-1080 cells grown in high glucose DMEM (Invitrogen, Ref # 11965084). Media was also supplemented with 10% fetal bovine serum (FBS, HiMedia, Cat # RM9952) and with non-essential amino acids (MEM-NEAA, Gibco, Ref# 11140-050) for MCF-7 cells. Cells were maintained at 37 °C at 5% CO_2_ and passaged at 70– 80% confluency using 0.25% trypsin-EDTA (HiMedia, Cat # TCL099). For all experiments, substrates were coated with 10 μg/cm^2^ collagen type-I from rat tail (Sigma, Cat # C3867) overnight at 4°C.

Stable ACTN4 knockdown cell lines were generated using MISSION^®^ Lentiviral Transduction Particles (Cat # SHCLNV, SigmaAldrich) as per standard manufacturer’s protocol. The sequences of the constructs are: CCGGCCTGTCACCAACCTGAACAATCTCGAGATTGTTCAGGTTGGTGACAGGTTTTTG (for shAC#1 cells) and CCGGGCCACACTATCGGACATCAAACTCGAGTTTGATGTCCGATAGTGTGGCTTTTTG (for shAC#2 cells). Stably transfected clones were selected by growing cells in the presence of puromycin (1.8 μg/ml for MDA-MB-231, and 1.5 μg/ml for HT-1080). Similar transfections carried out with an empty vector (Sigma, Cat # SHC001V), served as control. For fluorescence time-lapse studies, cells were transfected using transfection grade Polyethylenimine (Polysciences, Cat # 23966) or lipofectamine 3000 (Thermo, Cat # L3000001) as per standard protocol.

#### Preparation and characterization of Collagen gels

For fabricating collagen gels, required amounts of rat tail collagen type-1 (Corning, Ref# 354249) stock solution was mixed with 10× PBS and complete cell culture media, and the pH of the solution adjusted to 7.3 using 1 N NaOH solution. The collagen solutions were then incubated in the presence/absence of cells at 37 °C for 60-90 min for forming gels. For characterization of collagen gels, following snap freezing and fracturing of the hydrogels with an EM-grade blade, samples were mounted on the Cryo-unit (PP3000T, Quorum) of a JSM-7600F Cryo FEG-SEM. Samples were coated with a thin layer of platinum, and then images were obtained at desired magnification. Pore size was quantified using Fiji-ImageJ.

For gel compaction assay, 2 × 10^4^ cells were co-polymerized with 1.5 mg/ml rat tail collagen type-1 (Corning, Ref# 354249) on 48 well plates (200 μl/well) for 1 h at 37 °C and were cultured overnight in DMEM. After 24 h of culture, collagen gels were carefully released from the edges of the wells using sharp needles. 48 h after releasing the gels, the plate was imaged using a ChemiDoc^TM^ imaging system (BioRad). Gel compaction was quantified by measuring reduction in gel area calculated by the expression 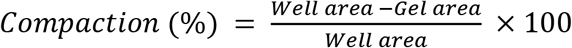

#### Single Cell Biophysics Measurements

For kymograph experiments, cells were sparsely seeded in collagen-coated 35mm dishes (Greiner) and live cell images were captured at 5 sec intervals for 15 min using an inverted phase-contrast microscope (Nikon Eclipse Ti, 20× objective). Protrusion and retraction rates were analyzed from obtained images using protocols as described elsewhere^21^. For live-cell images of myosin IIA dynamics, NMM IIA eGFP transfected cells were seeded on collagen-coated glass bottom dishes and images were captured at 5 sec intervals for 20 min at 63× magnification using a scanning probe confocal microscope (Zeiss, LSM 780). Kymographs were obtained by processing acquired images using Fiji-ImageJ. For probing focal adhesion dynamics, mCherry Paxillin transfected cells were seeded on collagen-coated glass bottom dishes and were imaged at 30 sec intervals for 30-45 min at 63× magnification using a scanning probe confocal microscope (Zeiss, LSM 780). Focal adhesion lifetime was analyzed from acquired normalized image sequences using protocols as described elsewhere^22^.

For 2D motility studies, cells were sparsely seeded (2000 cells/cm^2^) in collagen-coated 48-well plates. 12-14 h after seeding, motility videos were captured for 12 h at 15 minutes interval in a live cell imaging chamber (OKO Lab) using an inverted microscope (Olympus IX83). Cell trajectories and speed were measured from obtained videos using manual tracking plugin in Fiji-Image J.

For 3D cell motility studies, cells harvested from 70% confluent cell culture dishes were mixed with precursor collagen gel solutions at a density of 18,000 cells/ml such that the final concentration of the solution was 1.2 mg/ml for MDA-MB-231 and 1.5 mg/ml for HT-1080. 180 μl of this collagen-cell mixture was dispensed in each well of glutaraldehyde functionalized 48-well cell culture plates and incubated at 37 °C for gel formation. After addition of media on top of the gels, cells were cultured for another 16-18 h inside the collagen gels prior to imaging. For MMP inhibition experiments, media was supplemented with 15 μM GM6001. 3D motility videos were captured and analyzed the same way as described above for the 2D motility experiments. Cell embedded hydrogels were also imaged using confocal reflection microscopy (CRM) at 40× magnification using Scanning Probe Confocal Microscope (Zeiss, LSM 780).

For trans-well migration assays, 5 × 10^5^ cells were seeded on the upper chamber of 24 well plate cell culture inserts containing 3 μm pores (Merck, Cat # 353096) coated with rat-tail collagen I (Sigma, Cat # C3867). For creating a gradient, the upper chamber was filled with plain media and the lower chamber filled media was supplemented with 20% FBS. 24 h (for HT 1080) to 48 h (form MDA-MB-231) after seeding, cells were fixed with 4% PFA. To count cell numbers, fixed cells were stained with DAPI (10 minutes). After staining, membranes were cut using a scalpel and mounted on glass slides using mounting media. Confocal z-stack images of the fixed membranes were acquired at 20× and 63× magnification using Scanning Probe Confocal Microscope (Zeiss, LSM 780) at identical gain and exposure settings. Images analysis and cell counting at the top and bottom of the chamber was done using Image J software. Translocation efficiency (*η*) was calculated as 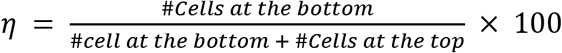. For trans-well migration rescue experiments, ACTN4 eGFP transfected knockdown cells were seeded in the trans-well chambers. GFP positive cells were counted at the top and bottom layer along with total cells and transfected cell ratio was calculated using the equation Normalized transfected cell ratio, 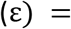 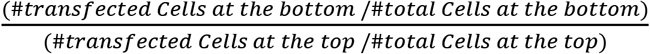.

Cell cortical stiffness was measured using an MFP3D Asylum AFM mounted on a Zeiss inverted microscope. Sparsely seeded cells were probed using a soft pyramidal cantilever with spring constant ~30 pN/nm (10 kHz, BL-TR400PB, Asylum research). Cells were probed slightly away from their center to minimize the effect of nuclear stiffness, also only the first 0.5 – 1 μm of indentation data was fitted to get the stiffness estimates. Obtained indentation data were fitted into the Hertz model to obtain estimates of cortical cell stiffness^23,24^.

#### Immunostaining

For immunostaining, cells were cultured on collagen-coated coverslips for 16 – 22 h prior to fixing with 4% paraformaldehyde. Fixed cells were permeabilized in 0.1% Triton X-100 for 8 – 12 min, blocked using 5% BSA 2 h at room temperature, and then incubated with primary antibodies (Anti-alpha Actinin 4 rabbit monoclonal antibody (Abcam, Cat # ab108198), anti-paxillin rabbit monoclonal antibody (Abcam, Cat # ab32084), Anti-non-muscle Myosin IIA rabbit polyclonal antibody (Abcam, Cat #ab75590), Anti-non-muscle Myosin IIB mouse monoclonal antibody (Abcam, Cat # ab684), anti-myosin light chain 2 (phosphor- Ser19) rabbit monoclonal antibody (Cat #3671, Cell Signaling)) diluted in PBS overnight at 4°C. Coverslips were then washed with PBS and were incubated for 1.5 h at room temperature (RT) with appropriate secondary antibodies diluted in PBS. Finally, nuclei were stained with Hoechst 33342 (Cat# B2261, Merck) for 5 min at room temperature and coverslips were mounted using mounting media. For, F-actin staining after secondary antibody incubation, Alexa-fluor 488 or Alexa-fluor 555 conjugated phalloidin (Thermo, Cat# A12379 and Cat# A12380) was added onto the coverslips for 2 h at RT in dark condition. Mounted coverslips were imaged using a scanning probe confocal microscope (LSM 780, Zeiss) using 63× objective. Images were processed and analyzed using Fiji-ImageJ software. Filament number and length of was quantified from f-actin stained images using an open-source Filamentsensor software^25^.

#### Real time PCR and Western blotting

Total RNA was eluted from knockdown and control cells using the RNeasy^®^ Plus mini kit (Qiagen, Ref # 74134). 2 μg of total RNA was reverse transcribed using High Capacity cDNA Reverse Transcription Kit (AppliedBiosystems, Ref. # 4368814) and was amplified using PowerUp^TM^ SYBR Green Master Mix (AppliedBiosystems, Ref. # A25742) in a QuantStudio 5 (AppliedBiosystems). PCR data were analyzed using comparative Ct method and Cyclophilin A expression was used to normalize gene expression data. Primers used for RT-PCR are as follows: ACTN1-Forward: 5’-CAAAGATTGATCAGCTGGAG, ACTN1-Reverse: 5’-CTCTACCTCATTGATGGTCC, ACTN4-Forward: 5’-AGTATGACAAGCTGAGGAAG, ACTN4-Reverse: 5’-CTGAAAAGGCATGGTAGAAG, NMM IIA-Forward: 5’-GTGAAGAATGACAACTCCTC, NMM IIA-Reverse: 5’-GATAAGTCTCTCAATGTTGGCTC, NMM IIB-Forward: 5’-CACTGAGAAGAAGCTGAAAG, NMM IIB-Reverse: 5’-TCTGCTCTTTATACTGGTCC, Cyclophilin A-Forward: 5’-TGGGCCGCGTCTCCTTTGA, Cyclophilin A-Reverse: 5’-GGACTTGCCACCAGTGCCATTA.

Whole-cell lysates were prepared using RIPA buffer (Sigma, Cat # R0278) containing protease inhibitor cocktail (Sigma, Cat # P8340) and phosphatase inhibitor cocktail (Sigma, Cat # P0044), were then centrifuged at 14,000 rpm for 10 min and protein concentration of the collected supernatant was determined using Bradford reagent. SDS-PAGE was performed with an equal amount (20 – 30 μg) of protein per lane and then were transferred onto a nitrocellulose membrane (PALL Life Sciences, Cat # 66485). Following transfer, the membranes were blocked with 5% BSA prepared in TBST (10 mM Tris-HCl, pH 8.0 containing 150 mM NaCl and 0.1% Tween 20) for 1 h at room temperature and were then incubated overnight at 4 °C with following primary antibodies diluted in TBST: Anti-alpha Actinin 4 rabbit monoclonal antibody (Abcam, Cat # ab108198), Anti-Vimentin mouse monoclonal antibody (abcam, Cat # ab8978), anti-integrin β1 rabbit monoclonal antibody (Abcam, Cat # ab155145), GAPDH rabbit polyclonal antibody (Abcam, Cat # ab9485), anti-Lamin A/C (phospho S392) rabbit polyclonal antibody (Abcam, Cat # ab58528), anti-Lamin A/C (phosphor S22) rabbit polyclonal antibody (Cat# 2026, CST), anti-Lamin A/C mouse monoclonal antibody (Abcam, Cat # ab8984, Anti-non-muscle Myosin IIA rabbit polyclonal antibody (Abcam, Cat #ab75590), Anti-non-muscle Myosin IIB mouse monoclonal antibody (Abcam, Cat # ab684). Blots were then washed with TBST and were incubated with one of the following HRP-conjugated secondary antibodies for 1.5 h at room temperature: HRP-conjugated anti-rabbit IgG (Invitrogen), HRP-conjugated anti-mouse IgG (Invitrogen). Finally, membranes were washed three times with TBST and were developed using a chemiluminescent ECL kit (Pierce, Cat # 32106) and images were acquired using a ChemiDocTM imaging system (BioRad).

#### Protein-protein interaction detection using Co-immunoprecipitation assay (Co-IP) and Proximity ligation assay (PLA)

Co-IP was performed using Protein G Immunoprecipitation Kit (Merck, Cat # IP50) as per the manufacturer's protocol. Briefly, whole-cell lysates were incubated at 4 °C for 12 h with Anti-alpha Actinin 4 antibody (Abcam, Cat # ab108198) or Anti-non-muscle Myosin IIA antibody (Abcam, Cat #ab75590). 30 μl slurry of protein G-agarose beads were added to the immunocomplexes in the IP spin column and incubated for 6 h at 4 °C. After 6-7 times vigorous washing in the spin column using 1× IP buffer, pellets were eluted from the column after heating for 5 min at 95 °C with 1× Laemmli buffer. Eluted samples along with whole-cell lysates as a control were then subjected to SDS-PAGE and Western blot analysis using Anti-non-muscle Myosin IIA antibody, Anti-alpha Actinin 4 antibody and GAPDH antibody (Abcam, Cat # ab9485) as described earlier.

Duolink^®^ PLA Red Starter Kit (Merck, Cat # DUO92101), which allows for endogenous detection of protein interactions was used to detect ACTN4/NMM IIA interaction. The kit was used as per standard manufacturer’s protocol using rabbit anti-alpha-actinin 4 antibody (Abcam, Cat# ab108198) and anti-mouse non-muscle myosin IIA antibody (Novus, Cat# H00004627-M06). The PLA signals were visible as red fluorescent spots when imaged using a confocal microscope (Zeiss, LSM 780).

#### Statistical analysis

Data distribution was tested using the Kolmogorov-Smirnov normality test. Based on the outcome, either parametric or nonparametric statistical test was performed. For parametric data, statistical analysis was performed using one-way ANOVA and Tukey’s test was used to compare the means. Mann-Whitney test was performed for non-parametric data. Statistical analysis was performed using Origin 9.1 (OriginLab Corporation), with p <0:05 considered to be statistically significant.

### Computational Model

For studying the importance of ACTN4-mediated cell protrusive forces in regulating confined cell migration, a plane strain finite element (FE) model of confined migration was created in ABAQUS using an explicit formulation for resolving large deformations, similar to our recent publication ^26^. In this model, pore entry of a 10 μm diameter cell with a 5 μm diameter nucleus was simulated through a deformable matrix for varying pore sizes (*φ* = {3, 5, 8} μm) (Fig. 6A). The cell-matrix interface was assumed to be frictionless. The components of the cell including the cell cortex (0.5 μm thick) and the cytoplasm were modeled as Kelvin-Voigt viscoelastic elements with the viscoelastic character of each component represented in the form of normalized creep compliance (Supp. Fig. 7A). The nucleus was modeled as an elastic solid encircled with a viscoelastic 50 nm thick membrane mimicking the Lamin-rich nuclear periphery^6^. The surrounding tissue was also assumed to be viscoelastic in nature. A Poisson's ratio of ν = 0.3, typical of compressible biomaterials^6^, was chosen for all the viscoelastic elements and is consistent with experimental observations of cell migration through microchannels wherein cells get polarized in the direction of migration, but the accompanying lateral deformations are negligible^2,27^. All elastic material properties are listed in Supplementary Table 1.

In our model, pore entry was mediated by active protrusive forces exerted on 250 individual nodes at the cell front such that the magnitude of the maximum force generated by the cell (*F*_Pmax_) remained in the physiologically relevant range^28^ (Supp. Fig. 7B). *F*_Pmax_ was chosen to vary between 1.25 nN/μm and 6.25 nN/μm to mimic different levels of ACTN4. In line with our experimental observations, the cortical stiffness (*E*_*cortex*_) was also changed accordingly. The simulation was stopped once the nucleus entered the pore completely.

Different sections of the cell and the tissue were meshed such that a smaller element size was used at regions expected to undergo large deformations and/or coming in contact with the matrix. A total of 30551 bilinear plane strain CPE4R elements were used in the model with a minimum element dimension of 5 nm and a maximum of 20 μm. Mesh gradation of the entire system is shown in Fig. 6A. The nuclear membrane, 50 nm in thickness, had 10 elements in the through-thickness direction to mitigate the effects of excessive artificial bending stiffness.

## Results

### *α*-actinin-4 (ACTN4) expression is positively correlated with cancer cell motility

As per TCGA database^18–20^, higher ACTN4 expression in breast cancer cells is associated with lower overall survival as well as lower progression-free survival (Fig. 1A). Since cancer mortality is linked to cancer metastasis, to probe the role of ACTN4 in regulating invasiveness of breast cancer cells, ACTN4 levels and localization were probed in MCF-7, T47D, ZR-75 and MDA-MB-231 breast cancer cell lines. Of these, MCF-7 cells are tumorigenic but non-invasive, and MDA-MB-231 (hereafter MDA) cells are highly metastatic. In addition, invasive HT-1080 (hereafter HT) fibrosarcoma cells were also considered. MCF-7, T47D and ZR-75 cells, which possessed low ACTN4 levels (Fig. 1B), were also the least motile (Fig. 1C, D, Supp. Movie 1), with ACTN4 exhibiting a primarily cytoplasmic localization (Fig. 1E). In comparison, MDA and HT cells that possessed highest levels of ACTN4 were 5-6 fold more motile than the above cell types. In these invasive cells, ACTN4 was prominently localized at the cell periphery as evidenced by the clear enrichment at the cell periphery (Fig. 1E). Higher ACTN4 expression levels also correlated with higher expression of *β*1 integrins and the mesenchymal marker vimentin. Collectively, these results establish a clear correlation between ACTN4 levels and its membrane localization with cancer motility.

**Fig 1:**
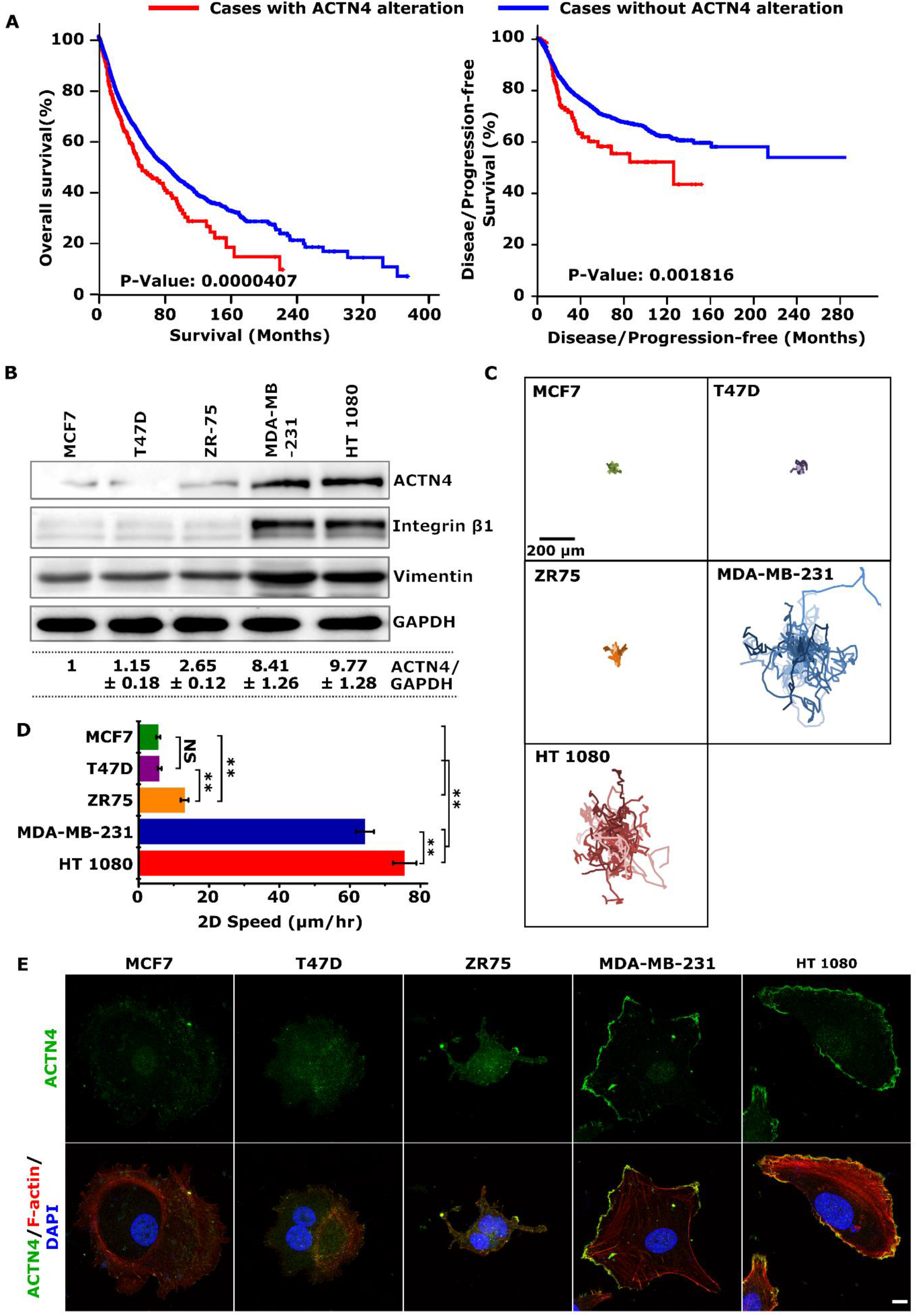
Actinin-4 expression correlates with cancer cell invasiveness. (A) Kaplan-Meier plots comparing overall survival (OS) and disease-free survival (DSF) in cases with or without ACTN4 alterations. The survival estimate was analyzed in http://www.cbioportal.org based on TCGA Pan-Cancer Atlas studies^18–20^. (B) Western blotting analysis of ACTN4, Integrin β1 and Vimentin expression in MCF-7, T47D, ZR-75, MDA-MB-231 and HT-1080 cells with GAPDH served as a loading control (*N* = 3, Values indicate relative fold change ± standard deviation). (C) Rose plots of migration trajectories of MCF-7, T47D, ZR-75, MDA-MB-231 and HT-1080 cells cultured on collagen-coated tissue culture plates. (D) Quantification of cell speed (*n* > 20 cells per condition across 2 independent experiments; ** p < 0.005, NS p > 0.05. Error bars represent standard errors of mean (± SEM).) (E) Maximum intensity projection images showing ACTN4 localization (green) in MCF-7, T47D, ZR-75, MDA-MB-231 and HT-1080 cells with merged image showing F-actin (red) stained by Alexa-fluor 555 conjugated phalloidin and nucleus (blue) stained by DAPI (Scale Bar = 10 μm).

### ACTN4 regulates cytoskeletal organization and focal adhesion dynamics

To probe the mechanisms of regulation of cancer invasiveness by ACTN4, stable knockdown cells (shAC#1, shAC#2) were established in MDA and HT cells (Figs. 2A, B). ACTN1 levels were unchanged in ACTN4 knockdown cells (Fig. 2C). Knockdown cells were found to be more elongated (i.e., less circularity) compared to control cells (shCTL), with an increase in spreading observed in MDA cells, but not in HT cells (Supp. Figs. 1A, B). However, these changes were not associated with alterations in the expression of epithelial to mesenchymal (EMT) markers integrin β1 and vimentin (Figs. 2A, Supp. Fig. 1C). In line with its crosslinking function, ACTN4 knockdown led to a marked reduction in the number of stress fibers (Supp. Figs. 2A, B). Increase in the average filament length in knockdown cells can be attributed to the disappearance of shorter actin filaments from the cell interior (Supp. Fig. 2C). Consistent with the reduction in the number of F-actin filaments, cortical stiffness probed with AFM revealed that knockdown cells were softer than controls (Supp. Figs. 2D, E).

**Fig 2:**
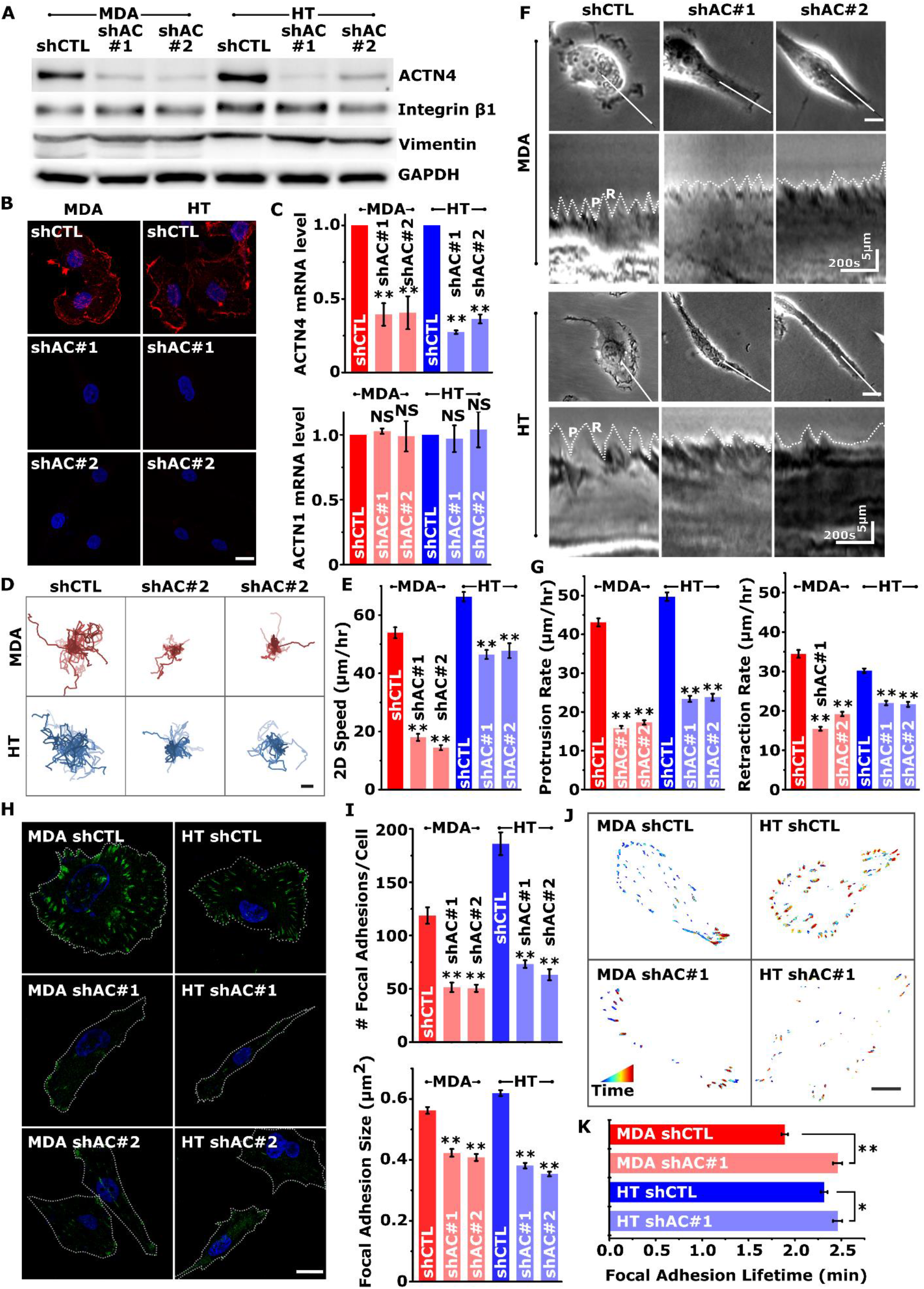
ACTN4 regulates cytoskeletal organization and adhesion dynamics. (A) Western blots showing actinin-4 (ACTN4), integrin β1 and vimentin levels in control (shCTL) and ACTN4 knockdown (shAC#1 and shAC#2) MDA-MB-231 (MDA) and HT 1080 (HT) cells with GAPDH serving as loading control. (B) ACTN-4 (red) immunostaining in control and knockdown cells, nucleus stained with DAPI (blue), scale bar = 10 μm. (C) Analysis of actinin-4 and actinin-1 (ACTN-1) mRNA transcripts using Real-time PCR. Total RNA harvested from 24 h culture of control and knockdown MDA and HT cells were subjected to quantitative real-time PCR analysis (n ≥ 2, ** p < 0.001; Error bars represent ± SEM). (D) Migration trajectories of control and ACTN4 knockdown cells cultured on collagen-coated tissue culture plates. Scale bar = 100 μm. (E) Quantification of cell motility of control and ACTN4 knockdown cells (*n* ≥ 20 cells per condition across 3 independent experiments; ** p < 0.001. Error bars represent ± SEM.) (F) Kymograph analysis of control and knockdown cells along the white solid lines. White dotted lines depict protrusion–retraction (P-R) cycles. Scale bars = 10 μm, unless otherwise indicated. (G) Analysis of protrusion/retraction rates in control and ACTN4 knockdown cells (*n* > 10 cells per condition across 3 independent experiments; ** p < 0.001. Error bars represent ± SEM.) (H) Representative paxillin stained (green) images of control and ACTN4 knockdown cells with nuclei stained with DAPI. Scale bar = 20 μm. (I) Quantification of average number and size of focal adhesions in control and ACTN4 knockdown cells (*n* = 10 - 30 cells per condition across 3 independent experiments; ** p < 0.001. Error bars represent ± SEM.) (J) Representative color-coded images depicting images of adhesions overlaid from multiple timepoints acquired over a period of 30-40 minutes. (K) Analysis of focal adhesion lifetime (*n* = 5 −8 cells were analysed per condition from 3 independent experiments; * p < 0.05, ** p < 0.001; Error bars represent ± SEM.)

Loss of ACTN4 led to reduced motility in both MDA and HT cells, with a more pronounced effect in MDA cells (Figs. 2D, E, Supp. Movie 2, 3). Kymograph analysis of leading edge protrusion dynamics revealed the presence of protrusion-retraction (P-R) cycles similar to that observed in other adherent cell types (Fig. 2F). Both the frequency of P-R cycles as well as the protrusion/retraction distances covered in each P-R cycle were reduced in knockdown cells, leading to reduction in both protrusion rate and retraction rate (Fig. 2G).

Transition from end of one cycle of retraction to initiation of another protrusion cycle coincides with formation of focal adhesions at the lamellipodia-lamellar interface^29^. Quantification of paxillin-positive focal adhesions revealed prominent reduction in both the number and size of focal adhesions in ACTN4 knockdown cells (Figs. 2H, I). Further characterization of paxillin turnover at focal adhesions revealed that adhesion lifetime was increased in knockdown cells (Figs. 2J, K, Supp. Movie 4, 5). Together, these results establish ACTN4 as a key molecule regulating cytoskeletal organization and adhesion dynamics, both of which are relevant to invasion.

### ACTN4 mediates cancer invasion by regulating actomyosin contractility

To next probe the importance of ACTN4 in regulating cancer invasiveness through 3D matrices, cells were embedded in 3D collagen gels with mean pore sizes of ~3 *μ*m for 1.2 mg/ml gels (used for MDA cells) and ~2 *μ*m for 1.5 mg/ml gels (used for HT cells) (Fig. 3A). While long ACTN4 enriched protrusions were observed in control cells (Supp. Fig. 3), drop in both the number of protrusions and the average protrusion length in knockdown cells (Figs. 3B, C) led to marked reduction in cell speed of ACTN4 knockdown cells (Figs. 3D, E, Supp. Movie 6, 7).

**Fig 3:**
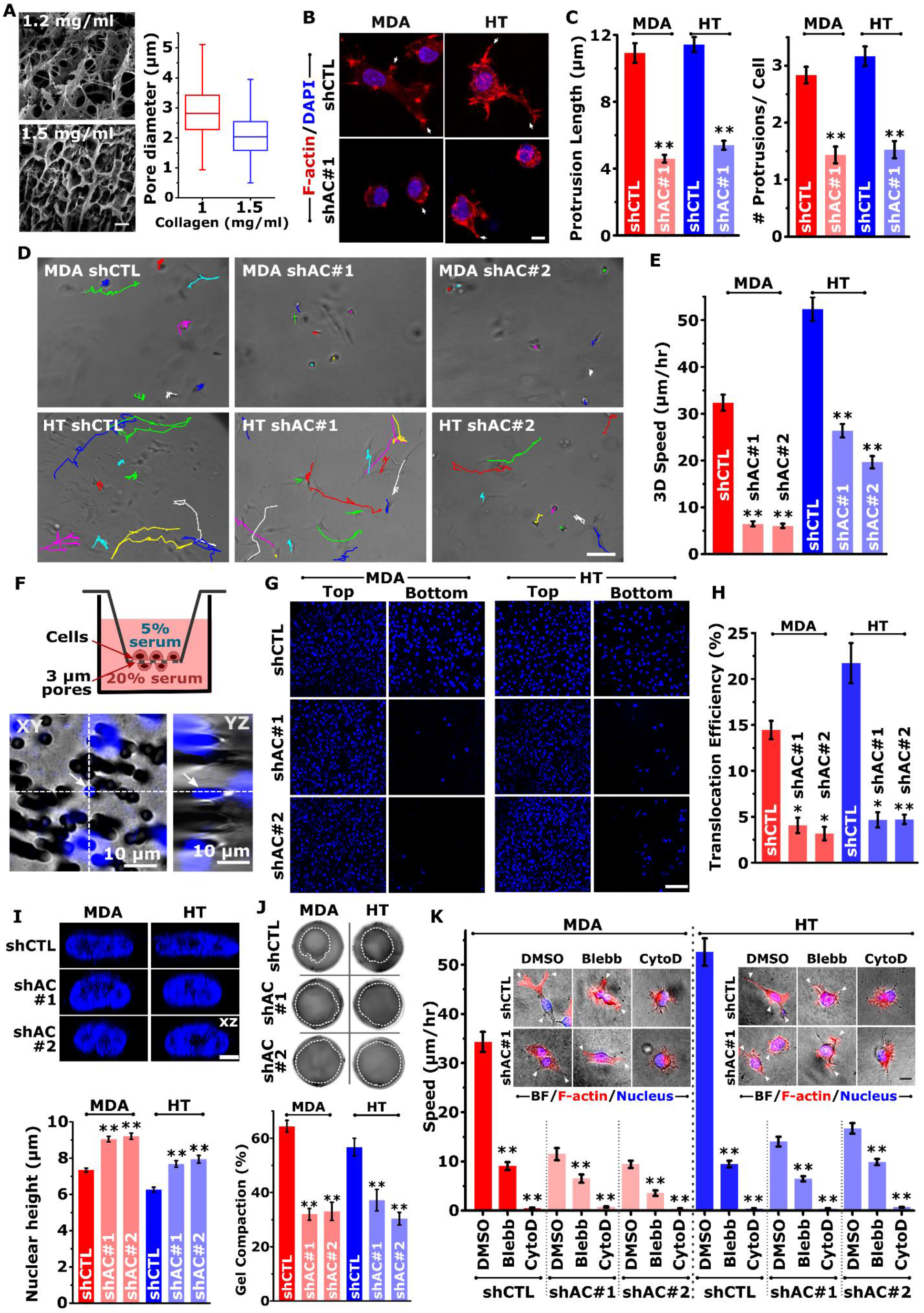
ACTN4 is essential for both proteolytic and non-proteolytic migration. (A) Representative Cryo FEG-SEM images of 1.2 mg/ml (top left) and 1.5 mg/ml (bottom left) 3D collagen gels and quantification of pore size (right). Scale bar = 2 μm. (B) Confocal images showing a greater number of protrusions (marked with arrows) in control cells compared to ACTN4 knockdown cells when cultured in 3D collagen. Merged images showing phalloidin stained F-actin (red) and DAPI stained nuclei (blue). Scale Bar = 10 μm. (C) Quantification of protrusion length and protrusion count per cell from acquired confocal images (*n* > 10 cells per condition across 3 independent experiments; ** p < 0.001 Error bars represent ± SEM). (D) Representative trajectories of control (shCTL) and ACTN4 knockdown (shAC#1 and shAC#2) MDA-MB-231 (in 1.2 mg/ml collagen gels) and HT-1080 (in 1.5 mg/ml collagen gels) cells migrating through 3D collagen gels. Scale bar = 100 μm. (E) Quantification of cell motility of control and ACTN4 knockdown cells in 3D collagen gel (*n* = 25 - 50 cells per condition across 3 independent experiments; ** p < 0.001). (F) Schematic of transwell migration assay through 3 μm sized pores (top) and representative XY and YZ plane orthogonal view of confocal microscopy image of a transmigrating nucleus. (G) Representative DAPI stained images of upper chamber (TOP) and lower chamber (BOTTOM) in control and ACTN4 knockdown MDA and HT cells (scale bar = 50 μm) and (H) its quantification (n > 700 cells per condition across 3 independent experiments; ** p < 0.001; Error bars represent ± SEM). (I) Representative side view images of DAPI-stained nuclei (top) of control and knockdown cells and quantitative analysis of nuclear height (bottom) (n = 15 – 30 nuclei per condition across 3 independent experiments; * p < 0.05, ** p < 0.001; Error bars represent ± Standard error of mean (± SEM)). Scale bar = 50 μm. (J) Representative images of cell laden gels (top) and quantification of gel compaction (bottom) (n = 3, ** p < 0.001; Error bars represent ± SEM). (K) Quantification of 3D cell motility of control and ACTN4 knockdown cells in the presence of DMSO (vehicle), Blebbistatin (Blebb) and Cytochalasin D (CytoD) (n = 40 - 125 cells per condition from 2 independent experiments; ** p < 0.001). Inset shows cell morphology after DMSO, Bleb and CytoD treatment with white arrows indicating protrusions. Monochrome brightfield images were merged with phalloidin stained F-actin (red) and DAPI stained nucleus (blue). Scale bar = 20 μm.

Nuclear deformation represents a rate limiting factor for migration in 3D matrices, particularly through sub-nuclear sized pores^8^. To probe the role of ACTN4 in mediating nuclear deformation, transwell migration experiments were performed using 3-micron size transwell pores (Fig. 3F). Comparison of the average number of nuclei at the top surface and bottom surface of the pores allowed us to compare the translocation efficiency of control and knockdown cells. In line with the above 3D motility findings, comparison of the fraction of cells that successfully transited through the transwell pores, i.e., translocation efficiency, revealed a dramatic drop in knockdown cells (Figs. 3G, H).

Lower translocation efficiency of ACTN4 knockdown cells was associated with an increase in average nuclear height of knockdown cells (Fig. 3I). Increased nuclear height of knockdown cells was associated with reduced contractility of knockdown cells, as assessed with gel compaction assay (Fig. 3J). Consistent with this, treatment of control cells with the myosin inhibitor blebbistatin (Blebb) led to drop in motility to levels comparable to that of knockdown cells (Fig. 3K, Supp. Fig. 4). While protrusions were still observed in Bleb-treated cells, inhibition of actin polymerization by cytochalasin D led to complete cell rounding and near complete loss of motility. Together, these observations suggest that ACTN4 regulates invasiveness by modulating protrusion dynamics and actomyosin contractility.

**Fig 4:**
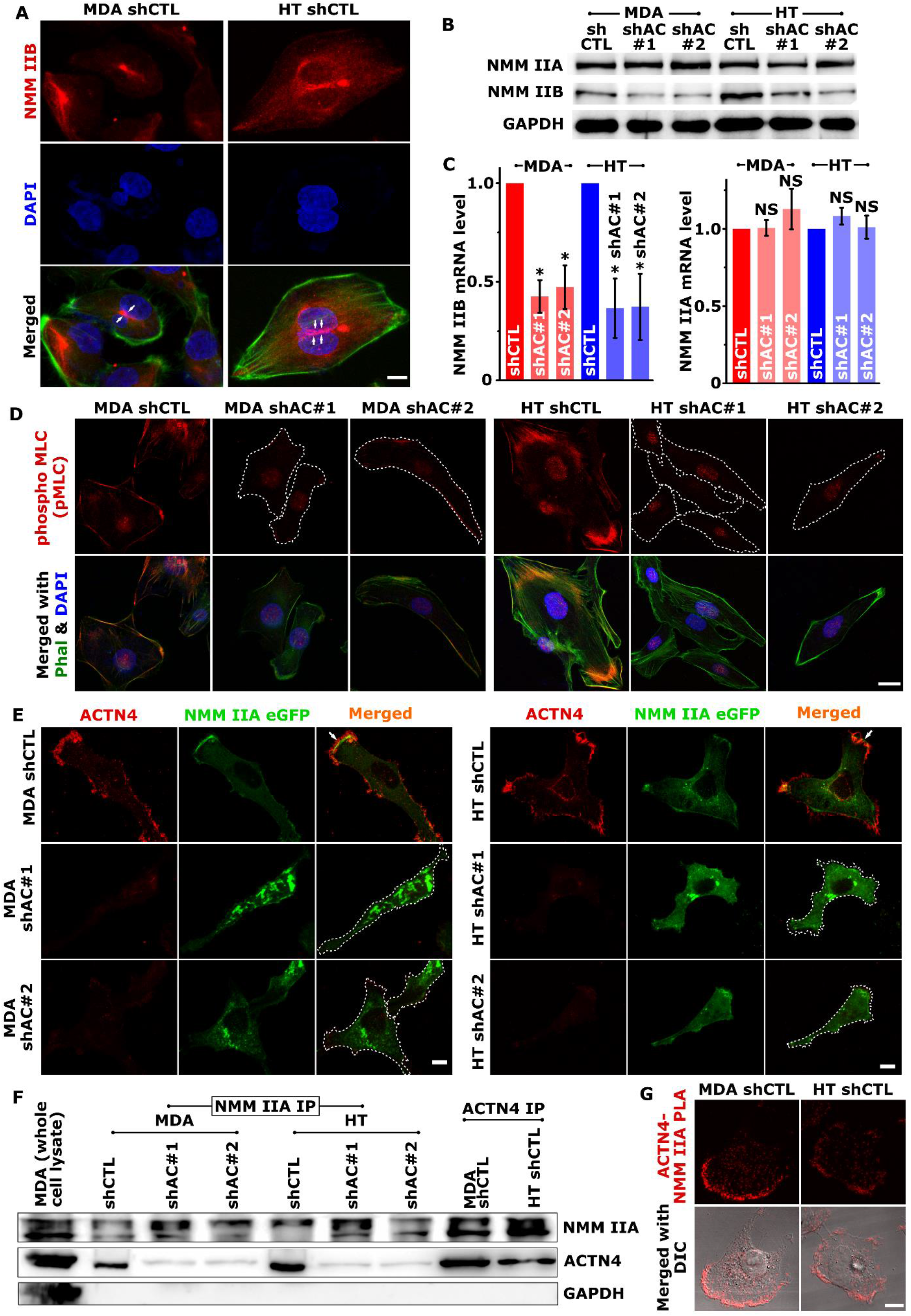
ACTN4 regulates myosin IIB expression and myosin IIA localization. (A) NMM IIB localization in control MDA and HT cells assessed using confocal microscopy. Representative maximum intensity projection (MIP) images showing NMM IIB (red) and nuclei (blue) staining. Merged images were generated by overlaying NMM IIB and DAPI staining with F-actin organization visualized with Alexa-fluor 488 conjugated phalloidin (green). White arrows indicate localization of NMM IIB at the sites of nuclear deformation (scale bars = 10 μm). (B) Western blotting analysis of non-muscle myosin IIA (NMM IIA) and non-muscle myosin IIB (NMM IIB) in control and knockdown cells with GAPDH serving as loading control (n ≥ 3). (C) Analysis of non-muscle myosin IIA (NMM IIA) and non-muscle myosin IIB (NMM IIB) mRNA transcripts using Real-time PCR. Total RNA harvested from 24 h old culture of control and ACTN4 knockdown cells were subjected to quantitative real-time PCR analysis (n = 3, * p < 0.05; Error bars represent ± SEM). (D) Representative maximum projection images of pMLC staining (red) in control (shCTL) and actinin-4 knockdown (shAC#1 and shAC#2) MDA-MB-231 (MDA) and HT 1080 (HT) cells. F-actin was stained with phalloidin (green) and nuclei were stained with DAPI (blue). Scale bar = 20 μm. (E) Control and ACTN4 knockdown MDA and HT cells were transfected with NMM IIA eGFP and were stained to visualize actinin-4 after fixing. Representative confocal MIP images show ACTN4 (red) and NMM IIA (green) localization in those cells (scale bars = 10 μm). (F) Co-immunoprecipitation study shows ACTN4-NMM IIA interaction. Lysates prepared from control and Knockdown cells were immunoprecipitated with either anti-NMM IIA antibody (NMM IIA IP) or anti-ACTN4 antibody (ACTN4 IP) and were subjected to western blot analysis using antibodies specific to ACTN4 and NMM IIA. MDA whole cell lysate was used as a positive control for the western blotting and GAPDH as a negative control for immunoprecipitation. (G) ACTN4-NMM IIA interaction detection using Duolink^®^ Proximal ligand assay. Images were acquired after the assay where red signal at the top panel indicates ACTN4– NMM IIA interaction and the merged differential interference contrast (DIC) image indicates the preferred location of the interaction is leading-edge (scale bar = 10 μm).

### ACTN4 regulates myosin IIB expression and myosin IIA localization

Efficiency of nuclear translocation through pores is dictated by physical properties of the nucleus including size and stiffness, as well as by the levels of non-muscle myosin IIB (NMM IIB), which plays a key role in mediating nuclear deformation^30^. In line with the literature, NMM IIB was detected spanning the top of the nucleus in control cells (Fig. 4A), and was found to co-localize with the peri-nuclear actin network visualized by staining cells with ACTN4 and F-actin (Supp. Fig. 5). Since average nuclear height of knockdown cells was found to be greater (Fig. 4F) without any alteration in Lamin A/C levels and its phosphorylation (Supp. Figs. 6A, B), we checked if myosin levels and/or activity are perturbed in knockdown cells. Expression profiling of NMM IIA and IIB revealed dramatic reduction in NMM IIB levels both at transcriptional level and at protein level in knockdown cells (Figs. 4B, C, Supp. Fig. 6C). In comparison, NMM IIA expression remained unperturbed in knockdown cells.

**Fig 5:**
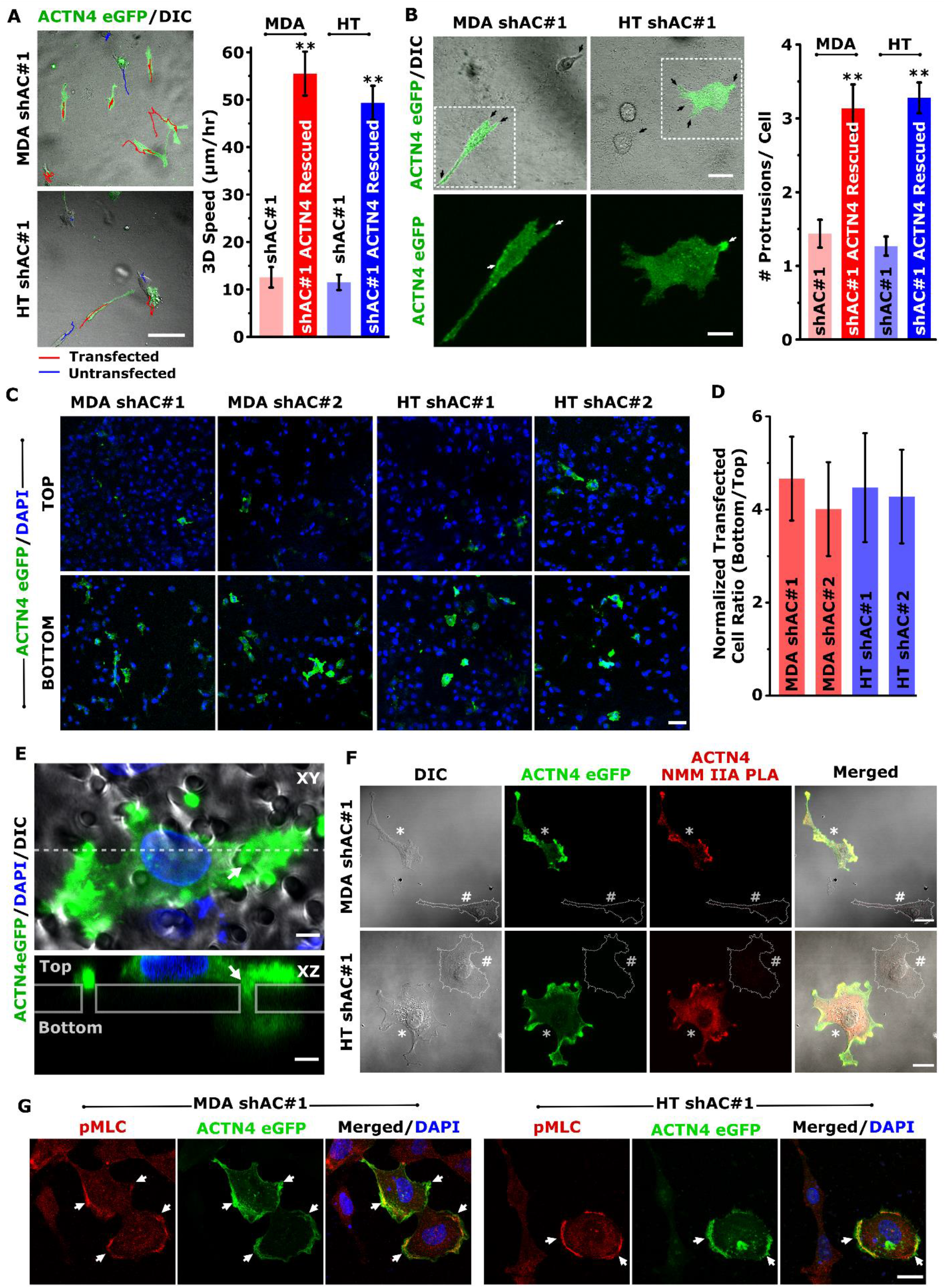
Ectopic ACTN4 expression in knockdown cells rescues pore migration and NMM IIA localization: (A) (Right) Representative trajectories of ACTN4 eGFP transfected (green) ACTN4 knockdown MDA-MB-231 (MDA shAC#1 in 1.2 mg/ml collagen gels) and HT-1080 (HT shAC#1 in 1.5 mg/ml collagen gels) cells migrating in 3D collagen gels. Red and blue lines indicating trajectories of transfected and untransfected cells, respectively. Scale bar = 100 μm. (Left) Quantification of cell motility of transfected and untransfected cells in 3D collagen gel (n ~ 40 cells per condition from 2 independent experiments; ** p < 0.001) (B) (Right) Representative images of protrusions (black arrows) in ACTN4 eGFP transfected knockdown cells. Bottom enlarged images show actinin-4 localization at protrusions (white arrows). Top Scale Bar = 30 μm, bottom scale bar = 15 μm. (Left) Quantification of protrusion count per cell from acquired confocal images (*n* = 30 – 56 cells per condition from 2 independent experiments; ** p < 0.001 Error bars represent ± SEM). (C, D) Representative images of the upper chamber (TOP) and lower chamber (BOTTOM) in transwell pore migration experiments with ACTN4 eGFP transfected ACTN4 knockdown cells and its quantification (n > 500 cells per condition from 2 independent experiments). Scale bar = 50 μm. (E) Representative XY and XZ plane orthogonal view (across dotted line in XY image) of ACTN4 eGFP transfected cell shows eGFP-ACTN4 localization (green) at sites of pore entry (white arrow). Scale bar = 5 μm. (F) ACTN4-NMM IIA interaction is rescued in knockdown cells when transfected with ACTN4 eGFP as detected using Duolink^®^ Proximal ligand assay. ACTN4 eGFP transfected cells (green; marked *) in acquired images show red signal indicating ACTN4–NMM IIA interaction while non-transfected cells (marked with #) lack any such signals. Scale bar = 20 μm. (G) Immunostaining shows prominent pMLC (red) co-localization with ACTN4 in the ACTN4 eGFP transfected (green) cells. Nuclei were stained with DAPI. Scale bar = 10 μm.

**Fig 6:**
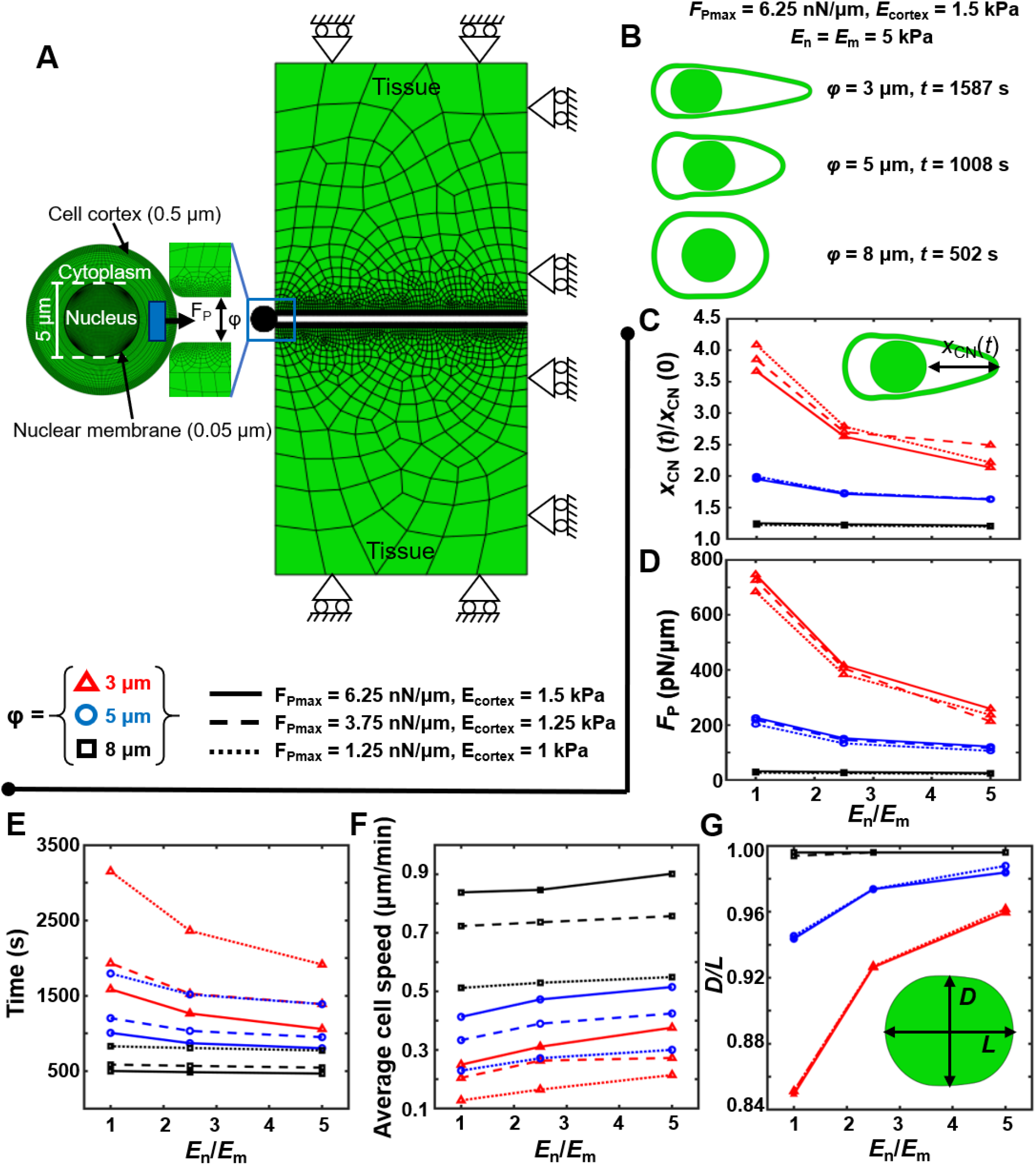
Model-based estimation of protrusion forces, cell speed and nuclear deformation during confined migration. A) Finite element (FE) plane-strain model of cell migration through deformable matrices/tissues. Cell migration through a pore of diameter *φ* (= 3, 5, 8 *μ*m) is mediated by cell-generated protrusion force (*F*_*P*_) in a defined region at the leading edge (blue rectangle). *F*_*Pmax*_ corresponds to the maximum protrusion force exerted by the cell. Boundary conditions of the system are also shown. Three different combinations of (*F*_*P*_, *E*_cortex_) correspond to three different expression levels of ACTN4 in the cell. These are: *F*_*Pmax*_ = 6.25 *nN*/*μm* and *E*_*cortex*_ = 1.5 *kPa* (solid); *F*_*Pmax*_ = 3.75 *nN*/*μm* and *E*_*cortex*_ = 1.25 *kPa* (dashed); *F*_*Pmax*_ = 1.25 *nN*/*μm* and *E*_cortex_ = 1 *kPa* (dotted). (B) Representative deformed cell profiles upon entry into pores of different sizes corresponding to *F*_*Pmax*_ and *E*_cortex_ of 6.25 *nN*/*μm* and 1.5 *kPa* respectively. Nucleus stiffness (*E*_*n*_) and matrix stiffness (*E*_*m*_) were held constant at 5 kPa. (C-G) Quantification of normalized cellular elongation (C), predicted *F*_*P*_ to enter a pore (D), pore entry timescale (E), average cell speed (F) and nuclear circularity upon pore entry (G) as a function of the ratio between nucleus and matrix stiffness (*E*_*n*_/*E*_*m*_) for different pore sizes (*φ* = 3 *μm* (red triangles), 5 *μm* (blue circles), and 8 *μm* (black squares)) and different combinations of (*F*_*P*_, *E*_cortex_).

Though NMM IIA levels were unchanged in knockdown cells, there was a pronounced reduction in the activation of myosin light chain phosphorylation (pMLC) at the cell periphery (Fig. 4D). Tracking of localization dynamics in eGFP-NMM IIA transfected cells revealed prominent peripheral NMMIIA localization and its co-localization with ACTN4 in control cells. In contrast, NMM IIA exhibited a predominantly cytoplasmic localization in knockdown cells (Fig. 4E, Supp. Movie 8, 9). Co-immunoprecipitation (Co-IP) studies revealed a direct association between NMM IIA and ACTN4 in control cells, with proximity ligation assay (PLA) identifying the leading edge as the preferred location of association of these two molecules (Figs. 4F, G). Collectively, these results suggest that the invasive phenotype of cancer cells is maintained by ACTN4 through regulation of NMM IIB expression and via its association with NMM IIA and its stabilization at the leading edge.

### Ectopic ACTN4 expression in knockdown cells rescues pore migration and NMM IIA localization

To finally establish that the observed changes in knockdown cells are a direct consequence of ACTN4 knockdown, knockdown cells were transiently transfected with eGFP-ACTN4 yielding a transfection efficiency of ~20%. When embedded in 3D collagen gels, eGFP-ACTN4 transfected cells were found to migrate faster than knockdown cells and also exhibited increased protrusion formation (Figs. 5A, B, Supp. Movie 10). Comparison of the proportion of transfected cells between top and bottom of the pores revealed 4-5 fold enrichment of eGFP-ACTN4 transfected cells at the bottom of the pores (Figs. 5C, D), with eGFP-ACTN4 detected at sites of pore entry (Fig. 5E). Furthermore, in knockdown cells transfected with eGFP-ACTN4, peripheral localization of NMM IIA and its association with ACTN4 were re-established (Fig. 5F). Activated myosin (i.e., pMLC) was also found to co-localize prominently with ACTN4 at the cell periphery (Fig. 5G). Taken together, our results establish ACTN4 as a key regulator of cytoskeletal organization and nuclear deformability through modulation of NMM IIA localization and NMM IIB expression.

### Dynamics of pore migration: insights from simulations

Our results suggest that ACTN4 regulates invasion by regulating protrusion dynamics at the leading edge, and mediating nuclear deformation by regulating NMM IIB expression. To estimate ACTN4-dependent forces required for invasion, we have developed a finite element (FE) based computational model of confined cell migration accounting for physical properties of the cell and the nucleus (Refer to Methods for details). In our model, cell entry into a pore was mediated by protrusive forces *F*_*P*_ at the cell front, with higher *F*_*P*_ associated with higher ACTN4 levels and increased protrusive force generation (Fig. 6A). The region of generation of *F*_*P*_ is representative of localization of NMM IIA at the cell front (in the direction of migration, depicted in blue in Fig. 6A) andits association with ACTN4. The NMM IIB-mediated squeezing of the nucleus is encoded in the model as a consequence of FE mesh spring compression under external matrix-induced stresses during confined migration.

Simulations were performed for varying combinations of matrix stiffness (*E*_*m*_ = 1 – 5 *kPa*) and pore sizes (*φ* = 3 – 8 *μm*), while keeping nucleus stiffness (*E*_*n*_) constant at 5 kPa^24^. It was found that decrease in pore size led to an increase in cell elongation and time to migrate through the pore for *E*_*n*_ = *E*_*m*_ = 5 kPa (Fig. 6B). Cell elongation, measured as a ratio between the cortico-nuclear distance (*X*_*CN*_) at time instant '*t*’ to that at *t* = 0 (i. e., *X*_*CN*_(*t*)/*X*_*CN*_(0)) was found to increase with decrease in *F*_*Pmax*_ and *E*_*cortex*_, in concordance with our experimental observations (Fig. 6C). Correspondingly, the force *F*_*p*_ predicted by our model to be generated by a cell to enter a pore increased with increase in matrix stiffness and decrease in pore size (Fig. 6D). Since low ACTN4 density, NMM IIA and IIB expression correspond to low predicted cell generated forces, the time required by cells to migrate through confined pores subsequently increases (Fig. 6E). In simulations with very low *F*_*Pmax*_ denoting negligible ACTN4 levels, cells do not even migrate within the maximum computational time duration of 6000s, further confirming our experimental observation of stalling of cell migration.

It is interesting to note from our simulations that although the predicted *F*_*P*_ is considerably higher for 3 μm pores than 5 μm pores, the migration time required is similar for cells with low ACTN4 levels migrating through 5 μm pores (blue dotted line) and those with moderate ACTN4 levels migrating through 3 μm pores (red dashed line) (Fig. 6E). Average cell speeds increase with pore size, but decrease with decrease in ACTN4 levels and cortical stiffness (Fig. 6F). Average cell speeds in the range 6 – 54 μm/hr predicted by our model are comparable to experimental observations. Average cell speeds through large pores (*φ* = 8 *μm*) remain relatively unchanged whereas, small or moderate pores lead to an increase in cell speed with *E*_*n*_/*E*_*m*_.

The change in nuclear circularity predicted by our model during cell entry into a pore decreases maximally for *φ* = 3 *μm* and *E*_*n*_/*E*_*m*_ = 1 (Fig. 6G). A small increase in nuclear circularity is also observed for *φ* = 5 *μm* and *E*_*n*_/*E*_*m*_ = 5 for low versus normal ACTN4 levels which might be attributed to decrease in NMM IIB-mediated contractility in conjunction with ACTN4 depletion. Although our computational model makes some assumptions, however, these model results match well to our experimental observations. Taken together, they suggest that α-actinin-4 modulates cell migration by regulating the expression and localization of NMM IIA and IIB.

## Discussion

Our findings implicate ACTN4 as a master regulator of cancer invasiveness via its interaction with NMM IIA at the leading edge and regulation of NMM IIB expression (Fig. 7). Consistent with previous studies^31–33^, in both MDA-MB-231 and HT-1080 cells, ACTN4 knockdown led to reduction in the number of F-actin stress fibers and cortical softening. In contrast, in MDCK cells, ACTN4 knockdown has been associated with increased F-actin content. Increased stress fiber formation in these cells has been linked with increased F-actin association with tropomyosin, and is indicative of a competition between ACTN4 and tropomyosin for binding to F-actin^34^. The lack of F-actin stabilization in MDA-MB-231 cells may be attributed to absence or low expression of tropomyosin isoforms in these cells^35^.

**Figure 7:**
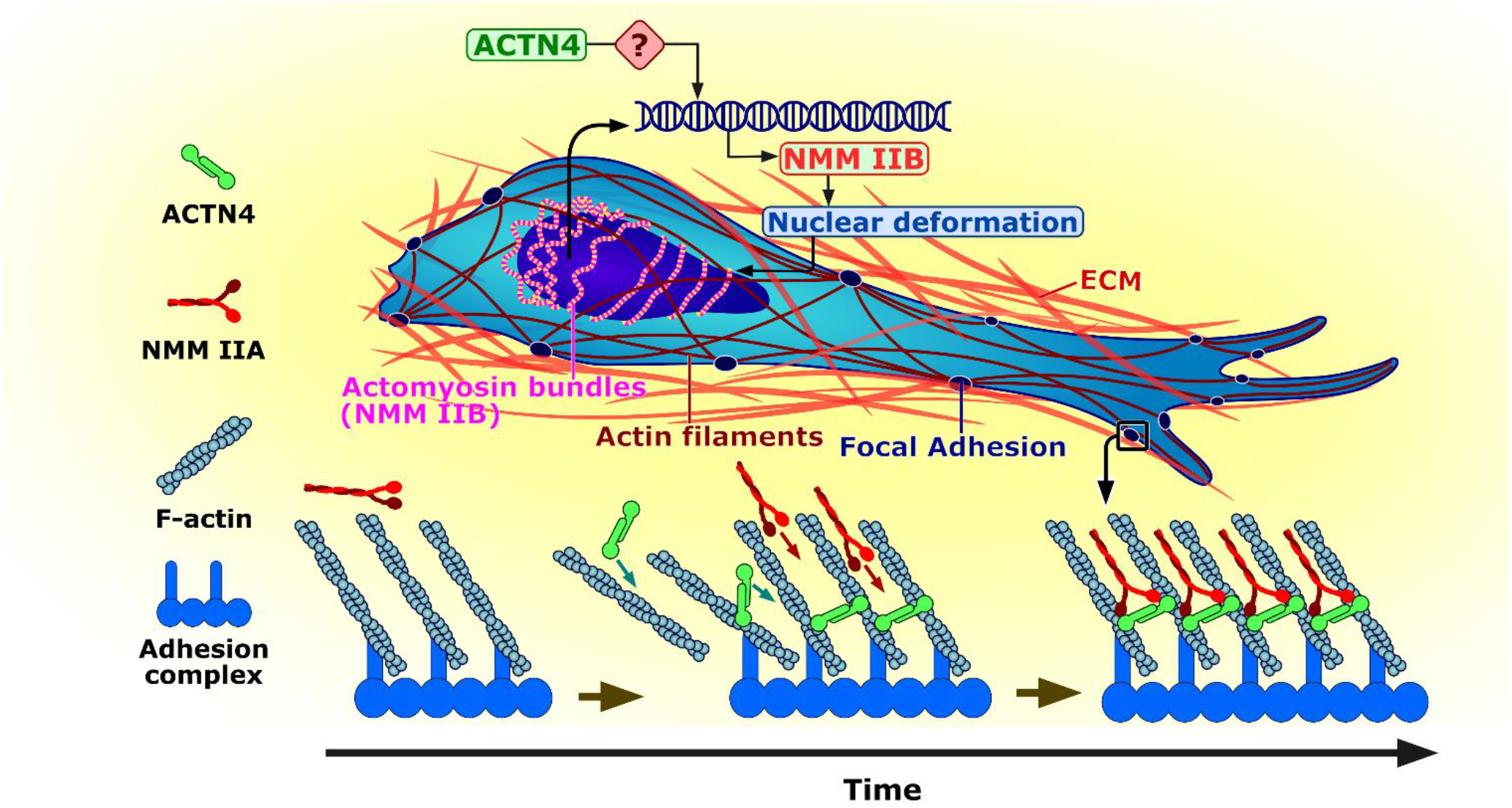
Schematic of regulation of cancer invasiveness by ACTN4. ACTN4-dependent retention of NMM IIA at the cell periphery and its phosphorylation drives focal adhesion maturation and cell migration. ACTN4 modulates nuclear deformability via transcriptional regulation of NMM IIB expression.

In addition to its actin crosslinking function, ACTN4 is increasingly known to regulate several aspects of cancer including cell proliferation^36,37^, epithelial to mesenchymal transition (EMT)^15^, and therapeutic resistance^14,38^. EMT is associated with transition from an epithelial cell-cell adhesion rich state to a mesenchymal state characterized by formation of cell-matrix adhesions. In both MDA-MB-231 and HT-1080 cells which are mesenchymal in nature, loss of paxillin-positive focal adhesions and increase in adhesion lifetime in ACTN4 knockdown suggests that ACTN4 is important for both adhesion formation and adhesion turnover. The inability of ACTN4 to bind zyxin in colorectal cancer cells has been implicated in preventing adhesion maturation thereby enabling faster turnover^39^. In melanoma cells that express high levels of ACTN4 and exhibit amoeboidal migration, ACTN4 knockdown led to transition from an amoeboidal morphology to a more mesenchymal state via increase in focal adhesion size^40^. These results thus suggest that ACTN4 levels dictate the mode of invasion, i.e., mesenchymal versus amoeboidal, by modulating the extent of cell-matrix adhesion.

Migrating cells consist of lamellipodia and lamella with the former composed of a branched actin network, and the latter enriched in more bundled actin^41,42^. While nascent focal adhesions are formed in the lammelipodial region in a force-independent manner, their maturation is brought about by coupling to actomyosin cytoskeleton to focal adhesions. Both *α*-actinin and NMM IIA have been shown to play prominent roles in focal adhesion maturation, with actin crosslinking function of *α*-actinin responsible for locally organizing the actin network, and NMM IIA-mediated contraction of the actin network mediating focal adhesion stabilization^43^. ACTN4 has been reported to participate in adhesion maturation by formation of stress fibers^44,45^. Parallelly, NMM IIA is also required for growth of focal adhesions, with adhesion maturation impaired in NMM IIA deficient cells^46,47^. Our studies suggest that in addition to locally organizing the actin network, ACTN4 plays a key role in retaining NMM IIA at the cell periphery and mediating NMM IIA phosphorylation. Thus, NMM IIA localization at anterior lamellar regions and maturation of focal adhesions by traction forces is directly regulated by ACTN4. Larger traction stresses at anterior focal adhesions in comparison to central adhesions may also be attributed to ACTN4/NMM IIA association at the cell periphery^48^. The increase in adhesion lifetime in ACTN4 knockdown cells may be associated with NMM IIA association with *α*-actinin-1 (ACTN1), which is also known to link F-actin to focal adhesions^49^, exhibits contrasting behaviour compared to ACTN4^50^, and has significantly higher actin binding affinity in comparison to ACTN4^51^.

When embedded in collagen matrices, MDA-MB and HT cells exhibit elongated morphologies with prominent localization of ACTN4 and F-actin along the protrusions. Loss of protrusions in ACTN4 knockdown cells suggests that leading edge protrusion is driven by polymerization of F-actin filaments stabilized by ACTN4 crosslinks. Under these conditions, physical resistance provided by the surrounding tissues may not only drive increased accumulation of ACTN4 given its mechanoresponsive nature, but also lead to increased actin binding thereby stabilizing the protrusions^52,53^. Our computational model predicts that during migration through larger pores (*φ* = 8 *μm*), protrusive forces (*F*_*P*_) required for pore entry and cell speeds were nearly constant over a wide range of *E*_*n*_/*E*_*m*_ values (1 − 5), and are consistent with the findings of statistical mechanics-based confined migration model^54^. While migrating through increasingly smaller pores, even for sustaining moderate cell motility (10 − 20 *μm*/*hr*), 10 − 20 fold higher protrusion forces (*F*_*P*_) are required, highlighting the importance of the mechanoaccumulative nature of ACTN4 required for mediating confined migration^52^. Reduction in gel compaction and peripheral myosin phosphorylation in ACTN4 knockdown cells, and drop in motility of Blebb-treated control cells to levels comparable to that of ACTN4 knockdown cells highlights the role of ACTN4 in driving confined migration via regulation of protrusion dynamics and actomyosin contractility. Our results predict protrusive forces, *F*_*P*_ scales with the degree of cell confinement (i.e. decreasing pore size *ϕ*) and with increasing ECM stiffness (i.e. decreasing *E*_*n*_/*E*_*m*_). This predictive relationship can be utilized to estimate protrusive force required by a cell to migrate through a pore in a matrix of given stiffness.

The nucleus, which is large and stiff, represents the rate limiting factor for 3D migration^55–58^. NMM IIB, which exhibits peri-nuclear localization, has been shown to be critical for mediating nuclear deformation and translocation by physical coupling the actomyosin cytoskeleton to the nucleus via the LINC complex^30,59,60^. Our observations of increase in nuclear height and concomitant loss of NMM IIB in ACTN4 knockdown cells are consistent with the above described function of NMM IIB^30,61^. Reduction in NMM IIB both at protein level and mRNA level are indicative of ACTN4 being a transcriptional regulator of NMM IIB. Thus, upregulation of NMM IIB in cells undergoing EMT^62^ maybe driven by ACTN4, which is an EMT inducer^15^. Though ACTN4 has been reported to translocate to the nucleus^63^, the mechanism(s) by which ACTN4 regulates NMM IIB expression remain(s) to be established. ACTN4 induces EMT by activating AKT and inducing expression of EMT associated transcription factors such as Snail and Slug^15^. Also, ACTN4 has been identified as a co-activator of several transcription factors including p65 subunit of NFkB^16^. However, the precise mechanisms of how ACTN4 regulates expression of multiple genes relevant to EMT including NMM IIB remains to be established and represents a future direction of research.

In conclusion, our results elucidate the mechanisms by which ACTN4 regulates cancer cell invasiveness. These include regulation of focal adhesion turnover, control over protrusion dynamics and contractility via its interactions with actomyosin network, and regulation of nuclear deformation by modulation of NMM IIB expression.

## Acknowledgments

The authors thank Dr. Alan Wells (Univ. of Pittsburg) for sharing the eGFP ACTN4 plasmid, Prof. Aurnab Ghose (IISER Pune) for sharing the mCherry-Paxillin plasmid, and Dr. Robert Adelstein (NIH, USA) for the GFP-NMM IIA plasmid. We acknowledge funding support from Department of Science and Technology (Grant # SR/ SO/BB-0016/2011, DST/SJF/LSA-01/2016-17) and IIT Bombay for providing Bio-AFM, Cryo FEG-SEM and Confocal Microscopy facilities.

## Author Contributions

AB and SS conceived the study. AB performed most of the experiments, with some help from AD and NS. AM performed the simulations. AB and SS wrote the manuscript.

## Supplementary Information

**Supplementary Table 1:** Material properties of system components.

| Component | Elastic Young's modulus, E (kPa) | Poisson's ratio ($\nu$) | Thickness or diameter ($\mu\text{m}$) |
| --- | --- | --- | --- |
| Cell cortex | 1 – 1.5 | 0.3 | 0.5 |
| Cytoplasm | 0.01 | 0.3 | 2 |
| Nuclear membrane | 5 | 0.3 | 0.05 |
| Nucleus | 5 | 0.3 | 4.9 |
| Tissue | 1 - 5 | 0.3 | 100 |

**Supplementary Figure 1:**
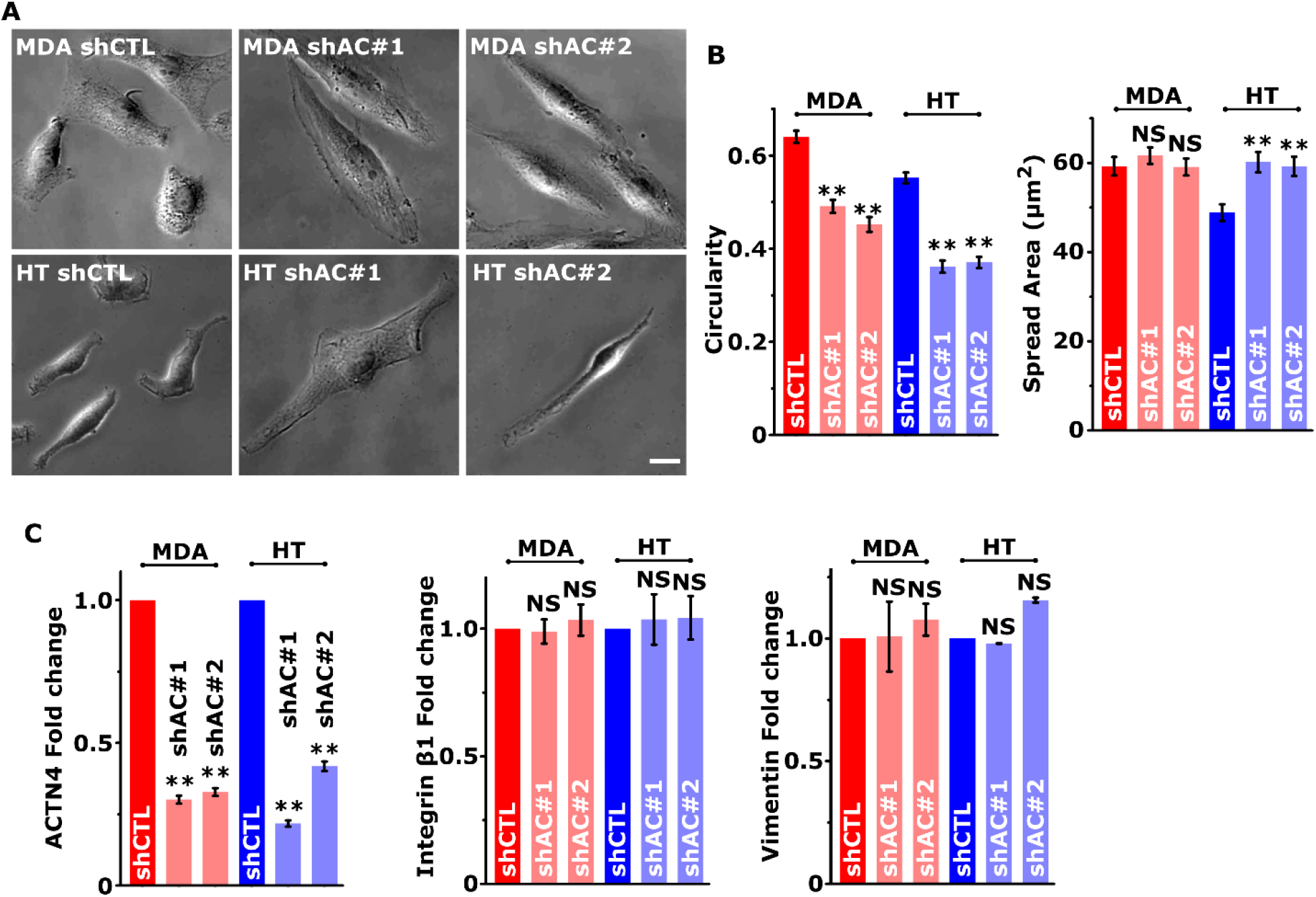
(A) Representative phase-contrast images of control (shCTL) and ACTN4 knockdown (shAC#1 and shAC#2) MDA-MB-231 (MDA) and HT 1080 (HT) cancer cells. Scale bar = 10 μm. (B) Quantification of cell spread area and circularity quantified from phase-contrast images. (n > 50 cells per condition from 3 independent experiments; ** p < 0.001, NS p > 0.05. Error bars represent ± SEM.) (C) Quantification of ACTN4, Integrin β1 and Vimentin western blots in with GAPDH serving as loading control (n ≥ 3 independent experiments; ** p < 0.001, NS p > 0.05. Error bars represent ± SEM).

**Supplementary Figure 2:**
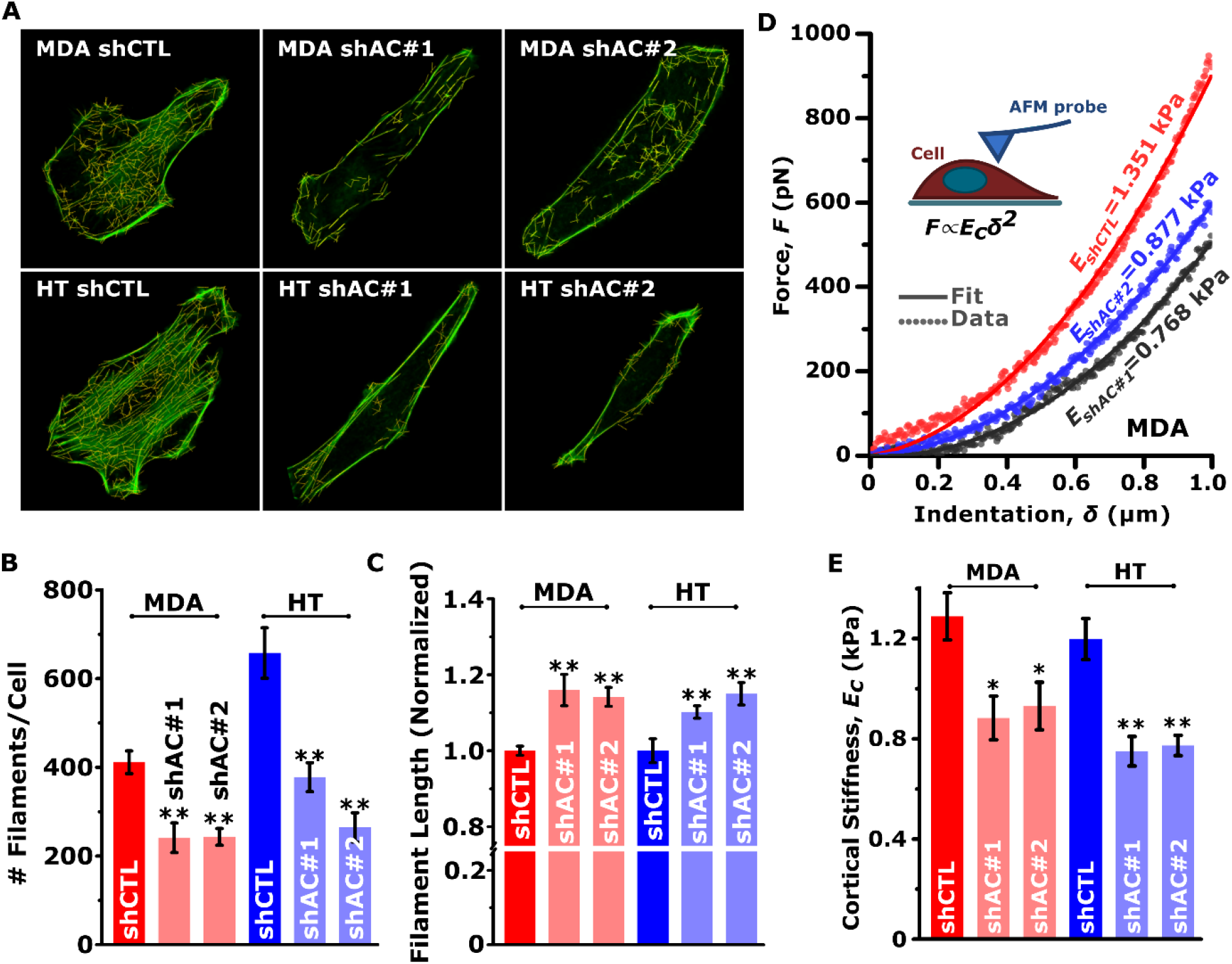
(A) Representative Alexa fluor-488 conjugated phalloidin stained F-actin images of control (shCTL) and ACTN4 knockdown (shAC#1, shAC#2) cells. Yellow lines represent F-actin stress fibers detected by image analysis software. Quantitative analysis of (B) number and (C) length of stress fibers in control and knockdown cells (*n* = 15 – 25 cells per condition across 3 independent experiments; ** p > 0.001; Error bars represent ± SEM.) Cortical stiffness of control and knockdown cells estimated using Atomic Force Microscope (AFM) by fitting < 1000 nm of indentation data using the Hertz model. (D) Representative force-indentation curves of control and ACTN4 knockdown MDA-MB-231 cells with inset showing schematic of the experiment setup. (E) Average cortical stiffness of control and ACTN4 knockdown cells (n > 150 cells per condition pooled from 3 independent experiments; ** p < 0.001, * p < 0.05. Error bars represent ± SEM).

**Supplementary Figure 3:**
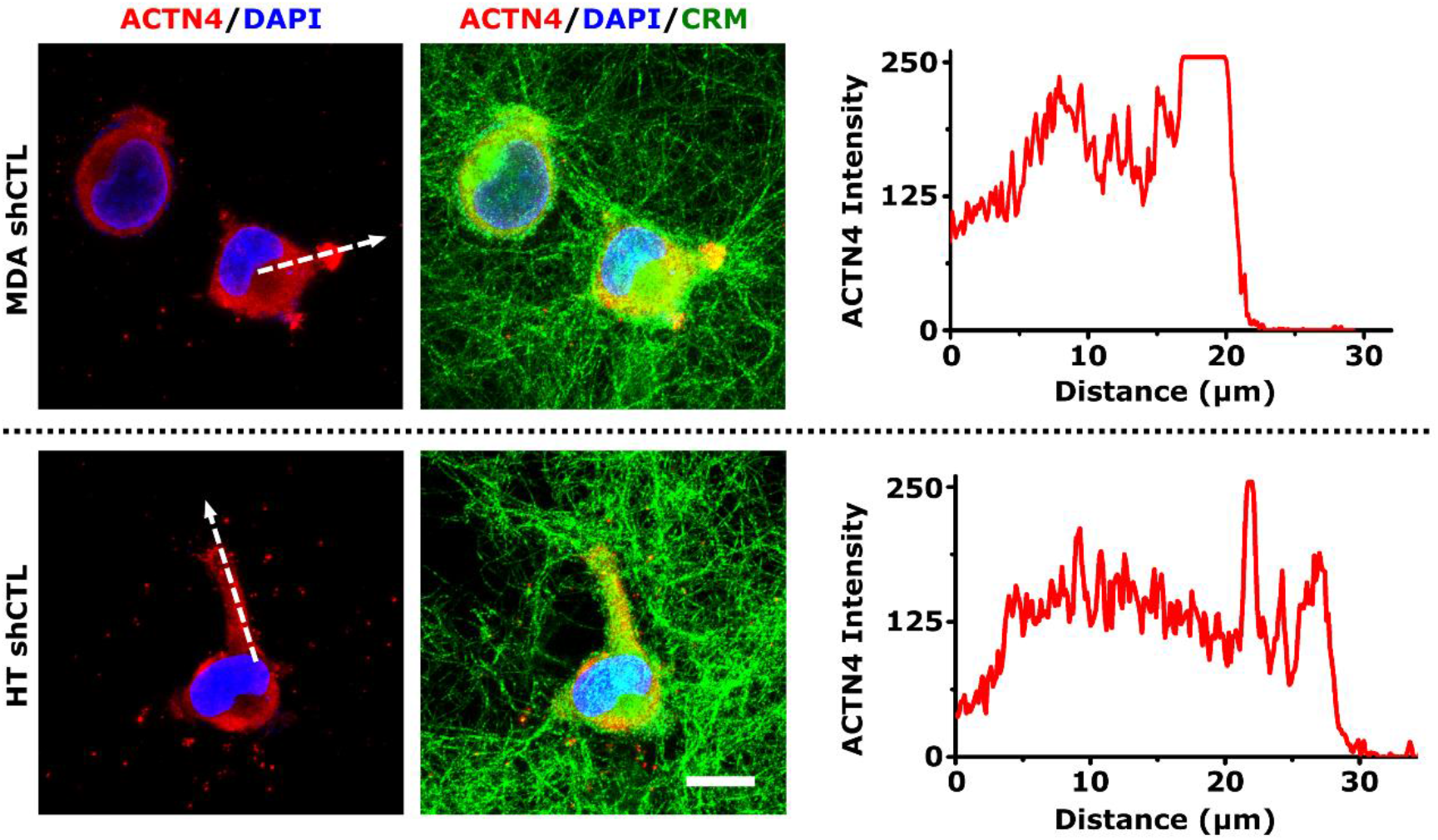
Immunostained confocal images showing ACTN4 localization (red) in cell protrusion of control MDA-MB-231 (MDA) and HT 1080 (HT) cells cultured in 3D collagen. Merged image is showing collagen fibers in green imaged using confocal reflection microscopy (CRM) and DAPI stained nuclei (blue). Scale Bar = 10 μm. The intensity plot is showing ACTN4 distribution in the cell protrusion along the white dotted line.

**Supplementary Figure 4:**
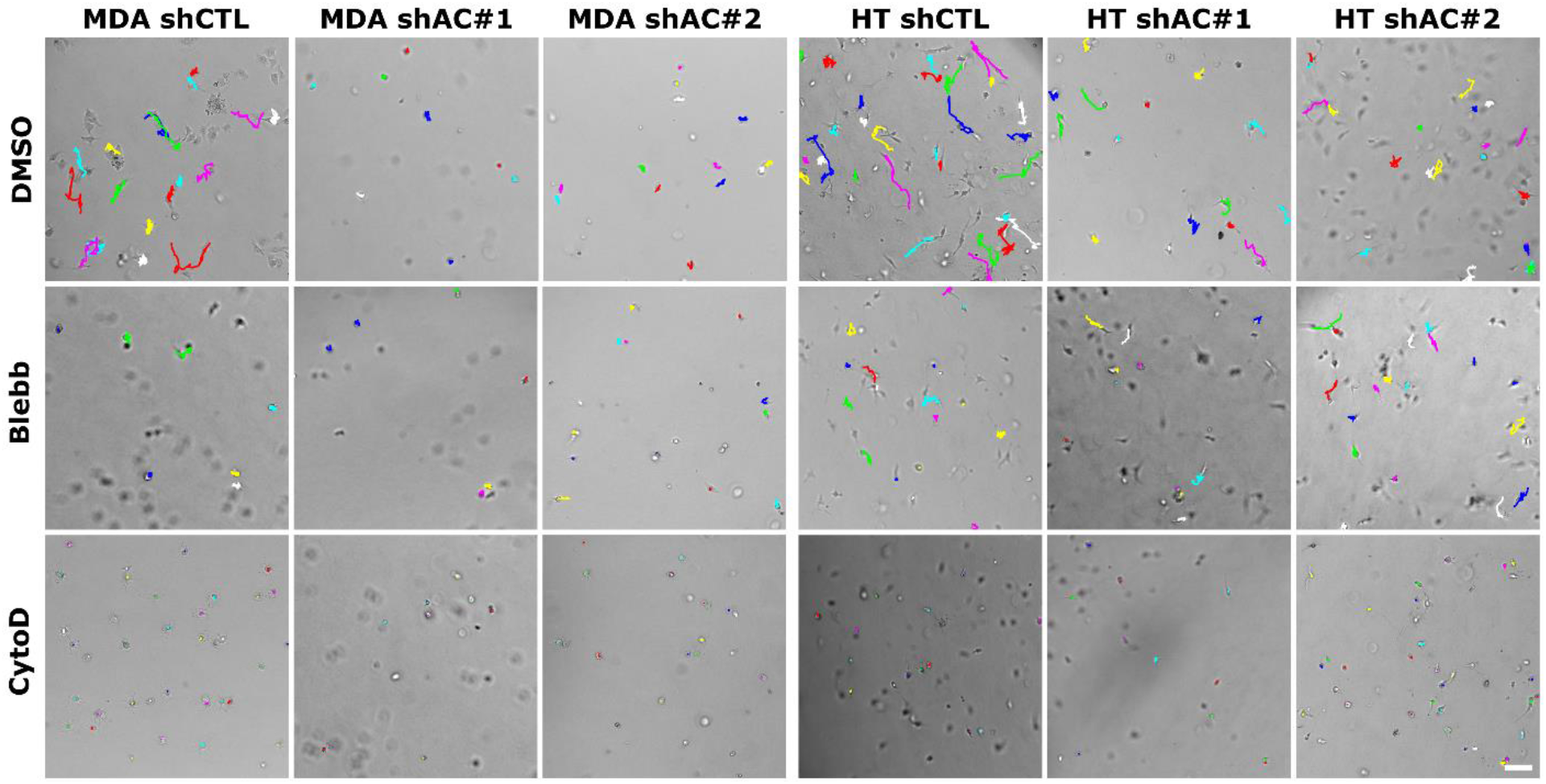
Representative trajectories of 3D cell motility of control (shCTL) and ACTN4 knockdown (shAC#1, shAC#2) cells in the presence of DMSO (vehicle), Blebbistatin (Blebb) and Cytochalasin D (CytoD).

**Supplementary Figure 5:**
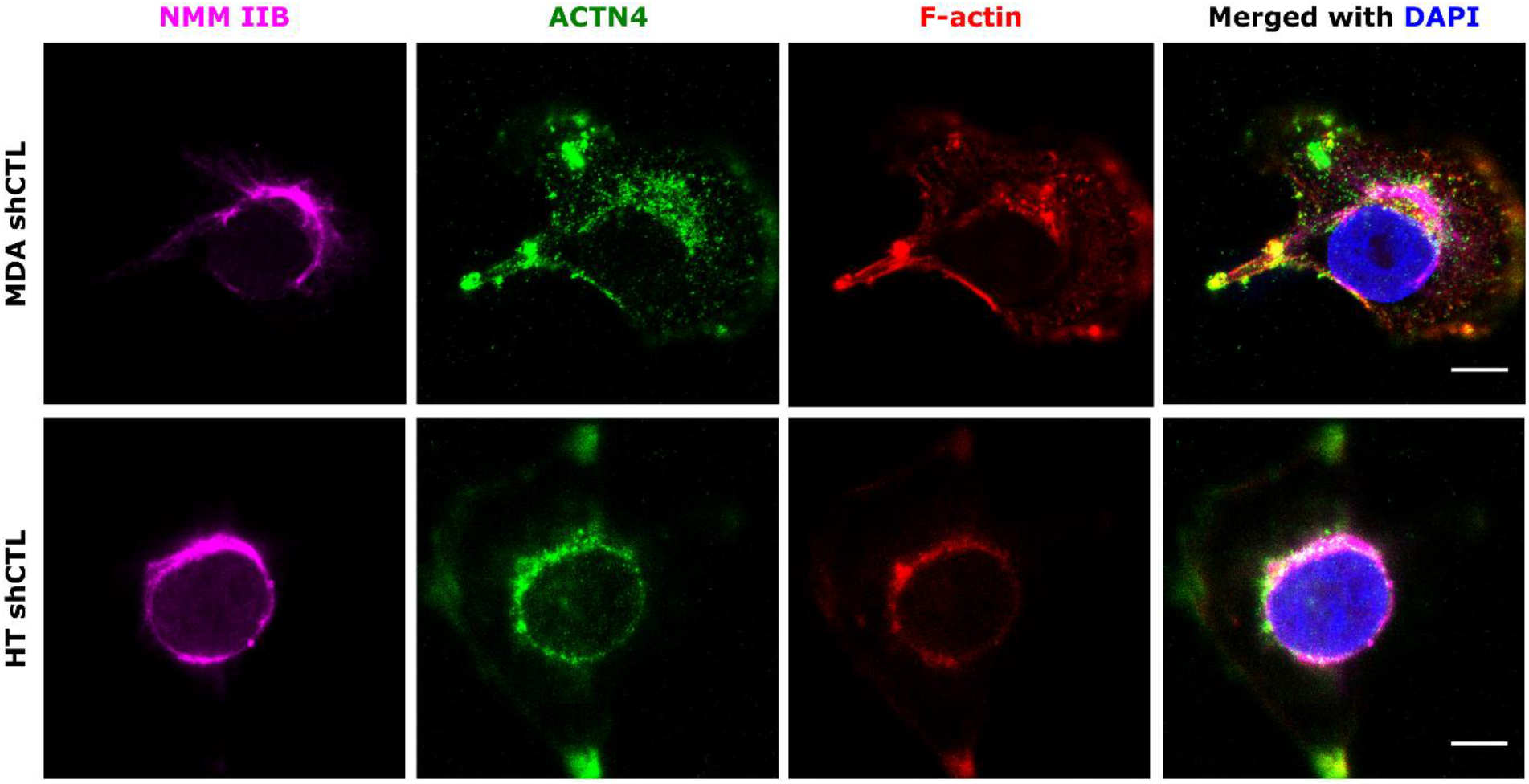
Perinuclear NMM IIB, Actinin4 (ACTN4) and F-actin localization in control MDA-MB-231 (MDA shCTL) and HT 1080 (HT shCTL) cells assessed using confocal microscopy. Representative images showing NMM IIB (red), ACTN4 (green), F-actin (red), and nuclei (blue). Cells were permeablized with 0.1% Triton X-100 before fixing to visualize perinuclear ACTN4 (scale bars = 10 μm).

**Supplementary Figure 6:**
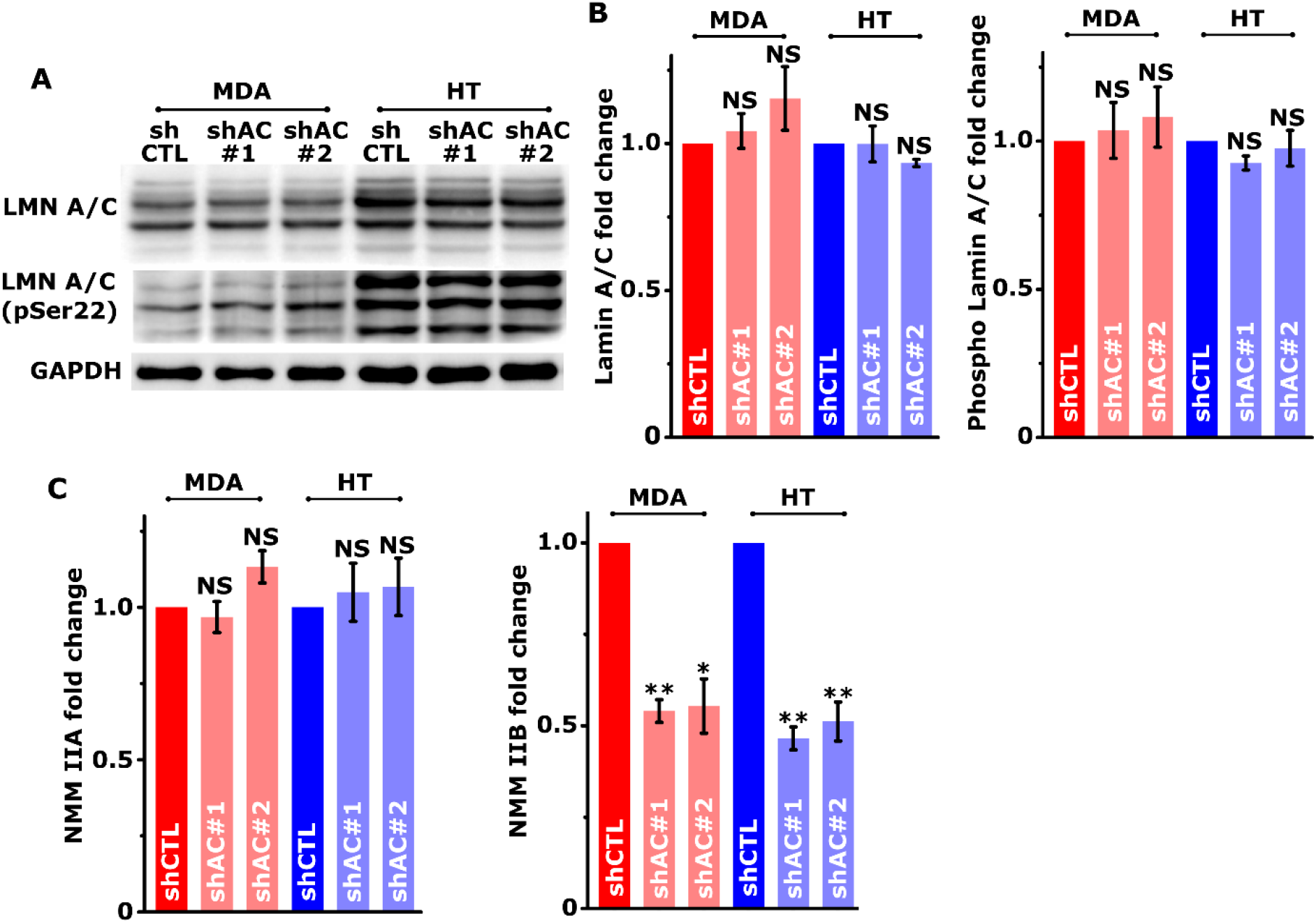
Western blotting analysis of Lamin A/C (LMN A/C) and phosphor Lamin A/C in control (shCTL) and knockdown (shAC#1 and shAC#2) MDA-MB-231 (MDA) and HT 1080 (HT) cells with GAPDH served as a loading control. (A) Representative images of western blots and (B) Quantification of western blots (n = 4, NS p > 0.05. Error bars represent ± SEM). (C) Quantification of NMM IIA and NMM IIB Western blots with GAPDH served as a loading control (n ≥ 3 independent experiments; ** p < 0.001, * p < 0.05, NS p > 0.05. Error bars represent ± SEM).

**Supplementary Figure 7:**
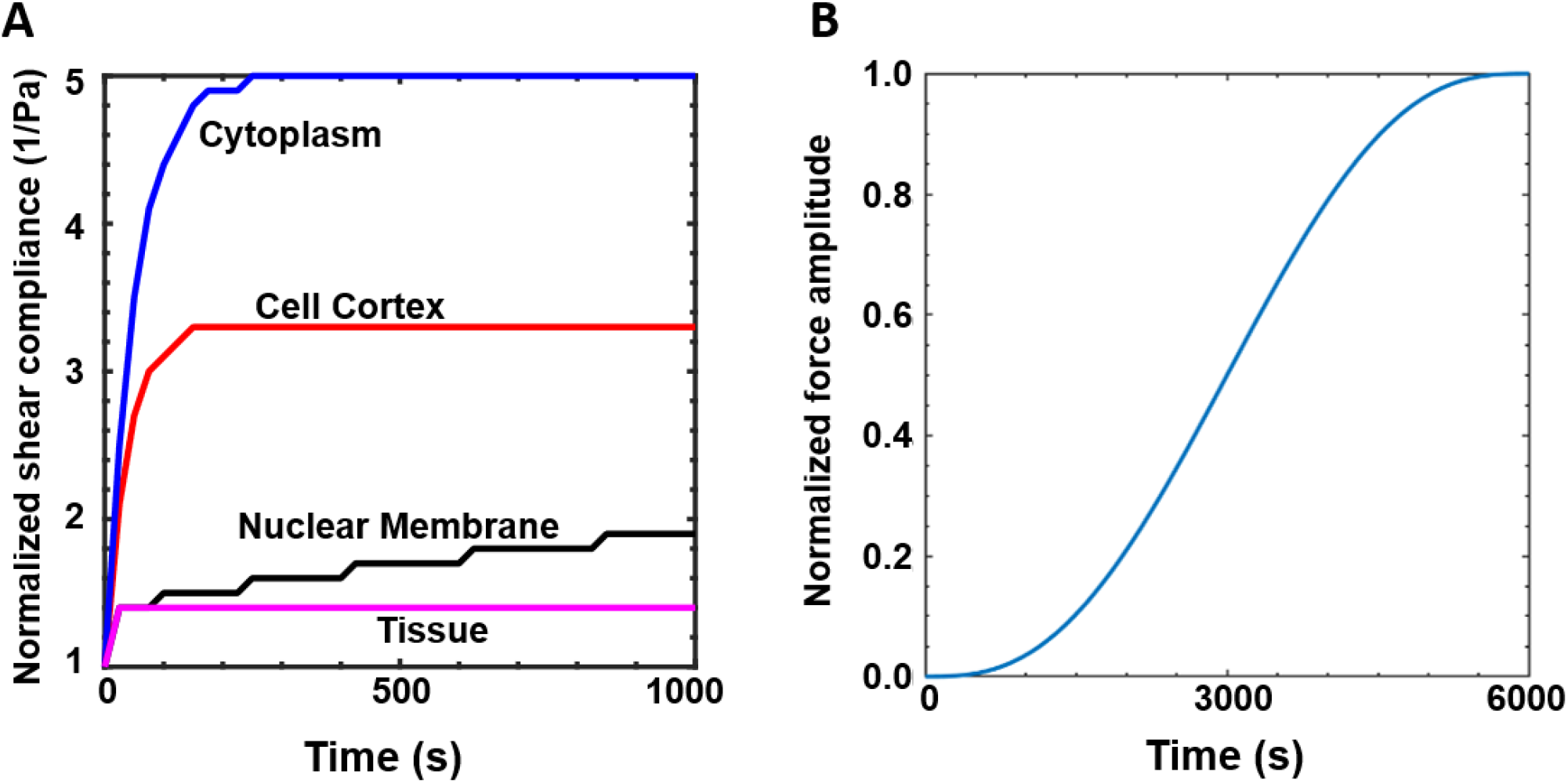
Computational model parameters. (A) Viscoelastic properties of the cytoplasm, cortex, nuclear membrane and extracellular matrix. (B) Rate of increase in protrusion force (F_Pmax_). For a maximum prescribed value of F_Pmax_, the input normalized force amplitude is ramped up as a sigmoid function in a time-dependent manner till a maximum time limit of 6000 s.

**Supplementary Movie Legends**

**Supp. Movie 1:** Migration trajectories of MCF-7, T47D, ZR-75, MDA-MB-231 and HT-1080 cells cultured on collagen-coated tissue culture plates. Lines depict trajectories of individual cells.

**Supp. Movie 2:** Migration trajectories of MDA-MB-231 control (MDA shCTL) and ACTN4 knockdown (MDA shAC#1, MDA shAC#2) cells cultured on collagen-coated tissue culture plates. Lines depict trajectories of individual cells.

**Supp. Movie 3:** Migration trajectories of HT 1080 control (HT shCTL) and ACTN4 knockdown (HT shAC#1, HT shAC#2) cells cultured on collagen-coated tissue culture plates. Lines depict trajectories of individual cells.

**Supp. Movie 4:** Movie showing tracking of Paxillin-mCherry focal adhesions in MDA-MB-231 control (MDA shCTL) and ACTN4 knockdown (MDA shAC#1) cells. The top panel shows normalized experiment acquired movies and the bottom panel shows adhesions outlined with different colors.

**Supp. Movie 5:** Movie showing tracking of Paxillin-mCherry focal adhesions in HT 1080 control (HT shCTL) and ACTN4 knockdown (HT shAC#1) cells. The top panel shows normalized experiment acquired movies and the bottom panel shows adhesions outlined with different colors.

**Supp. Movie 6:** Trajectories of MDA-MB-231 control (MDA shCTL) and ACTN4 knockdown (MDA shAC#1, MDA shAC#2) cells migrating through 1.2 mg/ml 3D collagen gels. Lines depict trajectories of individual cells.

**Supp. Movie 7:** Trajectories of HT 1080 control (HT shCTL) and ACTN4 knockdown (HT shAC#1, HT shAC#2) cells migrating through 1.5 mg/ml 3D collagen gels. Lines depict trajectories of individual cells. Lines depict trajectories of individual cells.

**Supp. Movie 8:** Non**-**muscle myosin IIA (NMM-IIA) dynamics in MDA-MB-231 control (MDA shCTL) and ACTN4 knockdown (MDA shAC#1, MDA shAC#2) cells after transiently transfecting the cells with GFP-NMM-IIA plasmids.

**Supp. Movie 9:** Non-muscle myosin IIA (NMM-IIA) dynamics in HT 1080 control (HT shCTL) and ACTN4 knockdown (HT shAC#1, HT shAC#2) cells after transiently transfecting the cells with GFP-NMM-IIA plasmids.

**Supp. Movie 10:** ACTN4 eGFP transfected (green) ACTN4 knockdown MDA-MB-231 (MDA shAC#1 in 1.2 mg/ml collagen gels) and HT-1080 (HT shAC#1 in 1.5 mg/ml collagen gels) cells migrating in 3D collagen gels. Lines depict trajectories of individual cells.

## Notes

### Competing Interest Statement

The authors have declared no competing interest.

